# INSIGHT: In Silico Drug Screening Platform using Interpretable Deep Learning Network

**DOI:** 10.1101/2025.04.21.649855

**Authors:** Xu Shi, Cyril Ramathal, Zoltan Dezso

**Affiliations:** AbbVie Bay Area, 1000 Gateway Boulevard, South San Francisco, CA 94080; AbbVie Inc., 1 North Waukegan Rd., North Chicago, IL 60064

## Abstract

The large-scale multiplexed drug screening platforms like PRISM and GDSC facilitate the screening of drug treatments over 1,000 cancer cell lines. The cancer cell lines are well characterized by multiomics screening in CCLE and DepMap, enabling the application of AI and machine learning techniques to study the association between drug sensitivity and the underlying molecular profiles. The large scale and variety of data modalities enabled us to build an interpretable deep learning framework, INSIGHT, integrating the multiomics data and the drug’s molecular structure to predict drug response. We trained our model on the PRISM screen for single treatments and on the DrugComb screen database for combination treatments. Our method enables the in-silico extension of current screens by predicting drug response in cancer cell lines not included in the screen, as well as the drug response to novel single agent or combination therapy by leveraging the drug’s molecular structure.

Furthermore, the deep learning framework was built to enable biological interpretation. The connections between the hidden layers of the neural network incorporate prior biological knowledge such as signaling pathways. This enables the model, in addition to predicting the drug sensitivity profiles, to prioritize the pathways predictive of drug response and identify pathways related to the mechanism of action (MOA) and potential off target effects of novel drugs.

The evaluation of our model using cross-validation on the PRISM and DrugComb dataset showed an improved performance compared to previously developed biologically informed deep learning methods and traditional state of the art machine learning methods like elastic-net and XGBoost. We illustrated with examples the value of incorporating biological knowledge into INSIGHT by relating the pathway activity of the predictive models to the MOA.

## INTRODUCTION

The traditional drug discovery process involves a series of costly and time-consuming development stages [1,2]. To expedite drug discovery efforts, high-throughput drug screening techniques have been developed to evaluate drug sensitivity across cancer cell lines. Platforms such as GDSC [3] and PRISM [4] have generated large scale datasets that measure the sensitivity of thousands of drugs across hundreds of cancer cell lines. Recently, DrugComb [5] built a database summarizing combination drug screens over cancer cell lines from multiple databases. The sensitivity measures across the databases are collected and standardized as combination sensitivity score (CSS). In addition, the Cancer Cell Line Encyclopedia (CCLE) [6] has generated comprehensive multiomics data, encompassing gene expression, mutations, and copy number variations for these cell lines. These extensive datasets present an unprecedented opportunity to develop machine learning integrative multi-omics models for in-silico drug screening.

In recent years, deep learning has shown great promise in the analysis of mutiomics data to predict drug response [7,8], cancer prognosis and the ability to uncover biomarkers [9–11]. Nevertheless, a significant challenge when employing traditional deep learning approaches is the aspect of interpretability. Artificial neurons, the fundamental units of these models, lack the inherent capacity to interpret the biological mechanisms underpinning their predictions. Most of these machine learning models are “black boxes” which are optimized for prediction accuracy, but provide no information about the biological mechanisms underlying the predicted outcomes [12]. To address the issue of limited interpretability, a novel paradigm has emerged in the form of biologically informed neural networks, wherein artificial neurons are encoding biological knowledge, such as Protein-Protein Interaction (PPI) networks and pathways [13]. Because the interconnections among these neurons are designed based on prior biological knowledge, the model can be interpreted by examining the activity of neurons which correspond to genes and pathways. DCell [14] implements a deep Visible Neural Networks (VNN) model designed to predict yeast growth in response to gene perturbation by the incorporation of the Gene Ontology (GO) network. Similarly, P-NET [15] develops a biologically informed network for predicting the disease state of prostate cancer patients. Another model, OntoVAE [16] introduces a biologically informed autoencoder architecture where the GO network is integrated into the decoder component. This framework shows its utility in inferring genetic perturbations and drug-induced effects by leveraging the pathway activity for analysis and interpretation.

For drug sensitivity prediction, Kuenzi et al. introduced DrugCell (DC) [17], an interpretable deep learning framework designed to model the area under the drug response curve (AUC) of drug sensitivity based on cell line mutations. This model consists of two key components: an interpretable neural network for embedding cell line mutations and an artificial neural network for embedding drug molecular structures. The interpretable neural network incorporates the GO to aid the interpretation of the drugs MOA through highlighting the relevant functions. In this study we extended the applicability of DC and improved its performance in several ways. First, the predictive model aims to capture the variances in drug sensitivity profiles across both samples and drugs. The variance across samples captures the differences of a drug’s sensitivity across cell lineages or mutations, while the variance across drugs reflects the differences in drug sensitivity profiles for specific cells or lineages. The DC model performs well in modeling the variance across drugs, due to the wide range of sensitivity profiles of the drugs, but it is not that effective in predicting the sensitivity across samples (Supplemental Figure S1). Furthermore, the current implementation of DC relies only on mutation data for modeling instead of utilizing multiomics integrative models, which have been demonstrated to enhance model performance [18]. Lastly, DC uses GO to capture the proteins’ biological function but does not include cancer related signaling pathways.

In this study, we developed an in-silico drug screening platform that builds on the DC framework while significantly expanding its functionality and overcoming some of the limitations of the model. First, we extended the model by including multiomics data to improve the performance of the model. Secondly, we built a platform with a novel two-level drug sensitivity model which captures both drug-level and sample-level variances, thereby enabling the differentiation of drugs based on their overall efficacy as well as prioritizing them for specific indications or molecular subtypes (Figure 1A). Finally, we encoded signaling pathways annotated by Reactome pathway database [19] in the model to better aid the biological interpretation. Our primary objective was to build a computational model that utilizes large scale cancer cell line screen (PRISM) data sets to predict drug sensitivity for unscreened cell lines based on their molecular profiles and novel compounds based on their molecular structure (Figure 1B).

**Figure 1.**
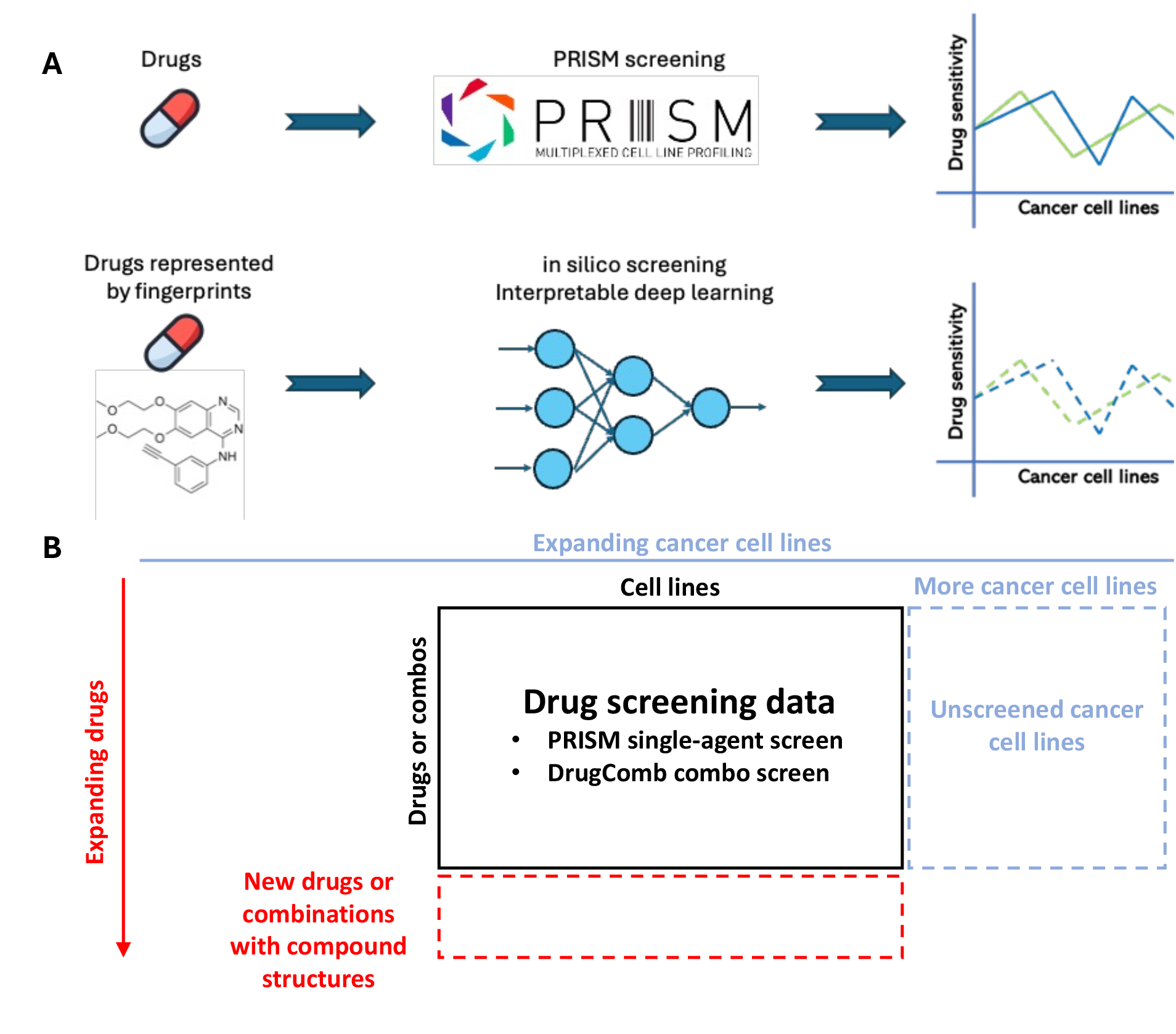
Drug sensitivity prediction overview. (A) Development of an in-silico screening platform utilizing an interpretable deep learning model to computationally evaluate drugs based on their fingerprints. (B) Applications of the drug screening model include evaluating more cancer cell lines and constructing sensitivity profiles for new drugs or drug combinations that have not been screened.

Many cancer cells, despite initially responding to treatment, can develop drug resistance through the activation of compensatory pathways. To achieve effective and lasting clinical responses, it became essential to identify drug combinations with the greatest potential for enhanced efficacy. Given the large number of potential drug combinations, it is unfeasible to test all of them experimentally. Therefore, there is a need for computational predictions to prioritize which combination to investigate experimentally. To this end, we extended the DC model functionality to include predictions for drug combinations based on the DrugComb database.

## RESULTS

### Description of the two-level drug sensitivity model

We developed a two-level deep learning framework to model the drug-level and sample-level variance separately (Figure 2A). The drug-level model was trained based on the drug-level sensitivity calculated as the average sensitivity of the drugs across all cancer cell lines. The sample-level model to better capture the sensitivity variance across samples was trained after first removing the variance across the drugs by scaling the sensitivity of each drug to zero. For drug sensitivity predictions, both models utilize drug fingerprints and multiomics data from the samples as input features. The final sensitivity prediction is obtained by summing the outputs generated by these two models (Figure 2A).

**Figure 2.**
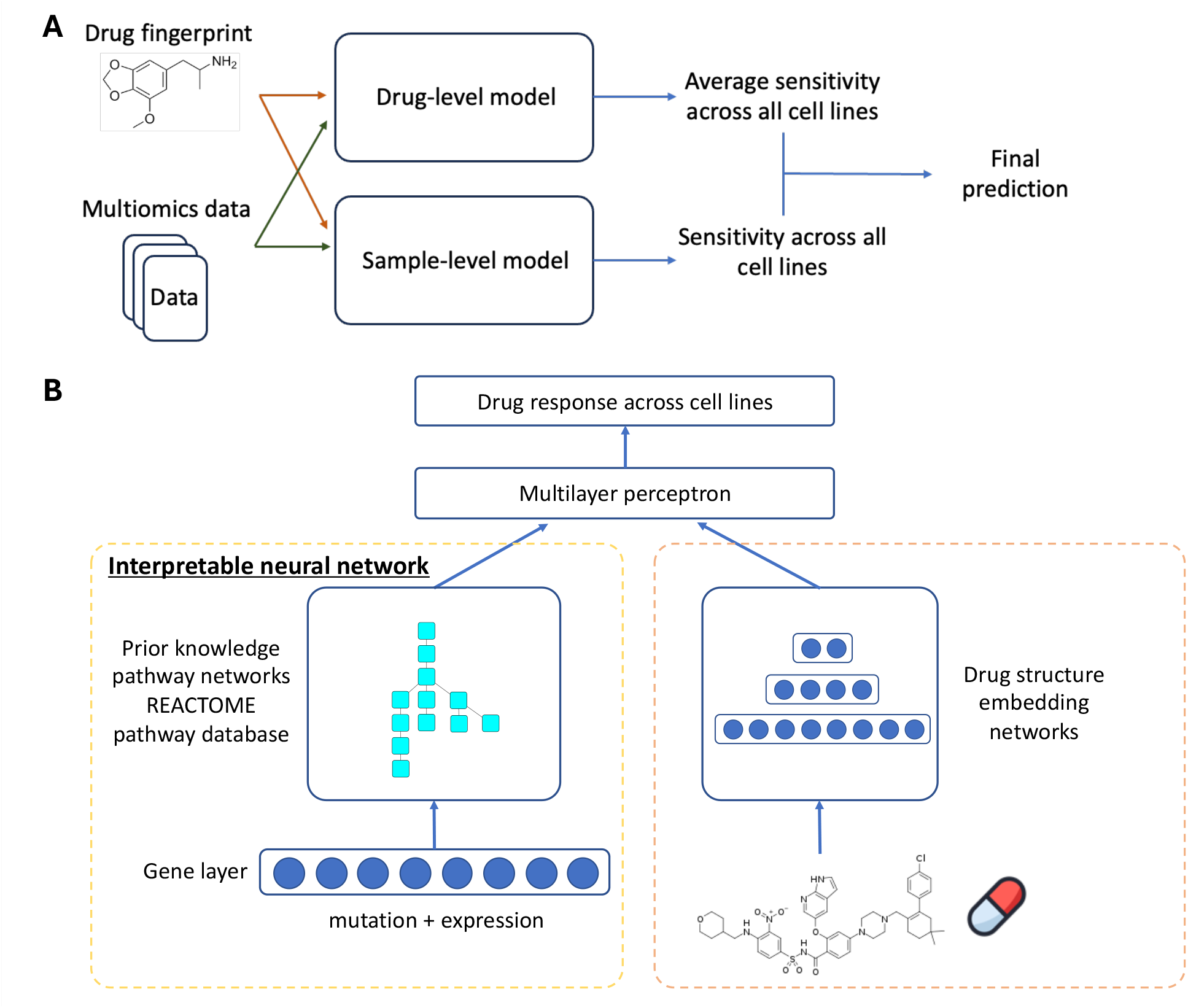
The framework of two-level drug sensitivity modeling. (A) Overview of the framework and (B) Detailed neural network structure

We designed the interpretable deep neural network framework inspired by DC as illustrated in Figure 2B. This framework comprises two components: one interpretable element that incorporates the pathway structure derived from established databases such as Reactome pathways, and another artificial neural network component that models the drug’s structural characteristics, represented in the form of a fingerprint [20].

### Performance evaluation of single-agent treatment on PRISM data

To assess the performance of our model we applied a 5-fold cross-validation based on the PRISM data. For the 5-fold cross-validation each drug dataset was divided into five folds and one-fold was held out for testing in each iteration. We compared our model’s performance against DC (version v1.0), and two state-of-the-art machine learning techniques, elastic-net (elnet, glmnet R package 4.1-8) and XGBoost (R package 1.7.6.1). The performance was evaluated by Pearson’s correlation between the measured PRISM sensitivity and the predicted values. Our model showed significant improvements in performance over the other methods tested (Figure 3A). The median performance of INSIGHT across all drugs demonstrated a substantial performance improvement, exceeding 38% in comparison to the second-best method DC (0.33 vs. 0.26). Figure 3B shows the comparison of performances of INSIGHT and DC across all drugs. The results indicate that INSIGHT and DC have comparable performance for a subset of drugs where both models perform well (correlation > 0.5). Interestingly, there is subset of drugs where DC failed to make good predictions, but INSIGHT was capable of building relatively well performing models. One reason for this increase in the performance may be due to the fact the DC model incorporates only mutation data, whereas INSIGHT is combining expression and mutation and for some drugs expression may be a better predictor of drug response. To test this, a feature selection analysis was done between the drug response and multiomics data using LASSO regression. Indeed, we found that more gene expression features were selected compared to mutations, and the expression coefficients were significantly higher compared to the coefficients of the mutations in the model, suggesting that the expression features have a strong contribution to the predictions (Supplemental Figure S2).

**Figure 3.**
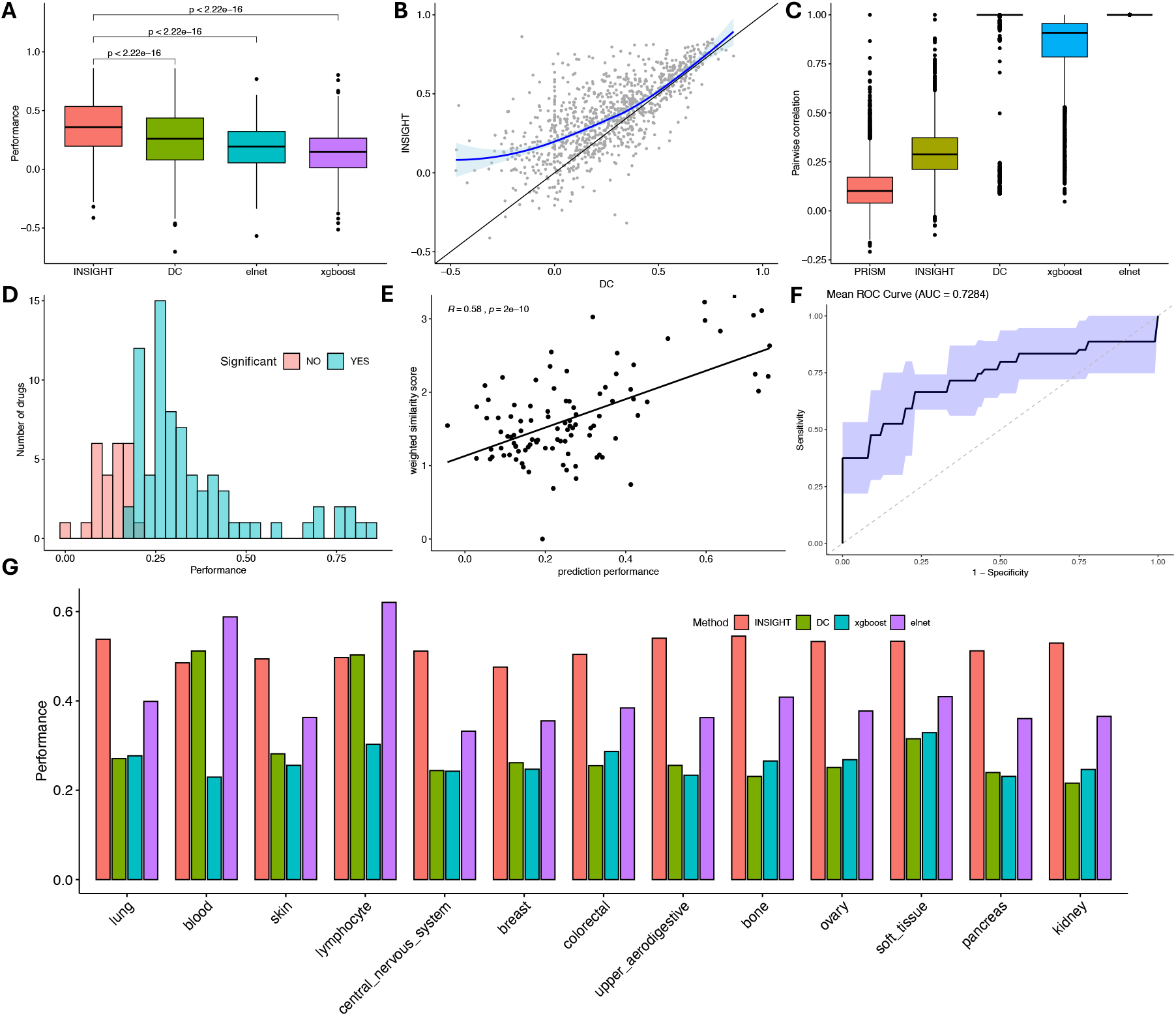
Performance on predicting drug sensitivity on PRISM datasets. (A) Performance of predicting sensitivity for independent samples. (B) Detailed comparison of performance between INSIGHT and DC. (C) Pairwise correlation of the prediction of test drugs over cancer cell lines. (D) Distribution of the performance of INSIGHT prediction on new drugs. A hypothesis test is done to label whether the correlation is significant. (E) Correlation between testing performance and weighted similarity score. (F) AUC of performance classifier test on 5-fold CV. (G) Performance on prioritize drugs for lineages with over 50 cell lines.

Besides predicting drug response to unscreened samples, the integration of drug fingerprints into our model enables the prediction of drug sensitivity for novel compounds not included in the training data. To assess the predictive performance for these novel drugs, we divided the 1,397 PRISM drugs into two groups. One group comprised 100 drugs, designated as the test set, while the remaining drugs were used for training the model. The molecular fingerprints of the structures of the drugs in the testing set and the multiomics data of all the cell lines were used as an input for the model to predict novel drug sensitivity. In this model the drug fingerprints encode the structural differences between the drugs, while the multiomics data captures the molecular profile variability among cell lines.

We first evaluated the ability of INSIGHT to differentiate the sensitivity of drugs by comparing it with DC, elnet and XGBoost. For elnet and XGBoost the multiomics data and the drug fingerprints were merged into the feature set of the predictive model.

To evaluate how well the models differentiate between drugs we computed the 4,950 pairwise correlation between the 100 test drugs and compared it to the observed pairwise correlation of the PRISM screen. Figure 3C shows the pairwise correlation distribution of INSIGHT as compared with the other methods. The elnet has all pairwise correlations of one, indicating that the linear method was unable to differentiate between any drug pairs. In fact, the elastic-net predictions for different drugs will only vary by a constant which explains that all pairwise correlation are one (Supplemental Methods S1). Although ensemble tree-based models like XGBoost can model non-linear relationships, the pairwise correlation was still very high with a median of 0.91 (Supplemental Figure S3). For the deep learning-based DC method, the pairwise correlation was likewise very high with a median very close to one (Supplemental Figure S4). The main reason is the lack of layers between the response layer and the embedding layers for pathways and drug fingerprints, making the model unable to differentiate between the sensitivity of drugs. Compared to these methods, INSIGHT achieved a significantly better pairwise correlation due to the inclusion of non-linearity layers, although it was still a little higher compared to the observed PRISM correlations. An example of correlations between two drugs based on the different methods are shown in Supplemental Figure S5.

The pairwise correlation analysis shows that current methods including DC, elnet and XGBoost were not suitable for predicting the drug sensitivity profiles over cancer cell lines. Therefore, we evaluated the performance of novel drug predictions only by INSIGHT. The performance of INSIGHT was evaluated by calculating the Pearson’s correlation between the predicted and observed PRISM sensitivity for each drug in the test set (Figure 3D, Supplemental Figure S6). A correlation significance test was performed to assess if the predicted sensitivity was significantly correlated with the measured PRISM sensitivity. There were 75 out of 100 drugs with significant correlation at FDR < 0.1 (Figure 3D).

Considering that some of the drug models did not pass the significance test for the drug sensitivity predictions, we evaluated our ability to identify our high confidence models based on the drugs fingerprint only. We hypothesized that a model would perform better if the novel drug had a structure which was similar to the structures of the drugs in the training set. To compare the drugs, we defined several structure similarity metrics based on fingerprint representations using both the Morgan and the infomax fingerprints (Methods). Since not all drugs can be modeled well by multiomics data, even if represented by drugs with similar structure in the training set, we developed a novel similarity metric score which combines as a weighted sum the structure similarity and fitting accuracy from the training data (see Methods). Figure 3E and Supplemental Figure S7 show that the prediction performance is associated with the structure similarity of test drugs and the drugs in the training set. Overall, we found that if the novel drug has a more similar structures in the training set, the INSIGHT model tends to have better performance. Among the three metrics, the weighted similarity had the highest correlation with model performance. Interestingly, we found that the prediction performance also correlated with the variance of sensitivity predictions over the cancer cell lines as shown in Supplemental Figure S7. The INSIGHT model will have better performance if the prediction values have larger variance, possibly reflecting the fact that both sensitive and resistant cell lines are well represented for model fitting. We finally trained a classifier combining the similarity measure and the variance of sensitivity via support vector machine (SVM). We evaluated the performance of the SVM classifier (R e1071 package 1.7-14) over the 100 test drugs by a 5-fold cross validation. Figure 3F shows the confidence level that a test drug’s performance can be classified has an average AUC of 0.7401 (R pROC package 1.18.5). The SVM classifier enables us to prioritize the novel drugs which can be considered for in-silico drug screening.

Although DC, elnet and XGBoost were not able to capture the variance across cell lines, it is still possible to apply them to predict the average sensitivity across multiple cell lines, which can be used to prioritize drugs for specific lineages. Leveraging the PRISM dataset, we assessed the model’s performance in prioritizing drugs across eight lineages with the highest number of cell lines in the PRISM data. For each lineage, we evaluated the relative sensitivity of the 100 drugs held out from the training set. Pearson correlation coefficient was used to measure the agreement between predicted and PRISM sensitivity, indicating the model’s ability to prioritize drugs based on their lineage sensitivity profiles. Figure 3G highlights that our model consistently outperforms existing models across various lineages.

### Performance evaluation on DrugComb database

In addition to single-agent, high-throughput combination screening enables the development of large-scale machine learning models to prioritize drug combination efficacy and reducing the large number of possible combinations to a smaller set suitable for experimental validations. One of the most comprehensive databases is DrugComb, integrating resources such as ALMANAC [21], ONEIL [22], FORCINA [23], and CLOUD [24]. After the preprocessing (refer to the Methods section for details), the input data comprises 11,412 drug pairs, including 280 drugs and 160 cell lines. Combination treatment sensitivity is quantified by the CSS, representing the inhibition percentage of the treatments.

To encode combination treatments in a fingerprint we followed the method utilized in DeepSynergy [25], where a combination’s fingerprint was represented by appending the fingerprints of the single drugs. Because the order of the appending does not impact performance, each prediction was duplicated with fingerprints added in both directions and the final predictions were averaged.

Similar to the single-treatment evaluation, we assessed the performance of predicting new cancer cell drug responses via 5-fold cross-validation on the DrugComb data. Many drug pairs have limited cell line screens in the DrugComb data, making them unsuitable for cross-validation. Thus, we selected 127 drug pairs with more than 50 cell lines to evaluate the performance of the predictions. Figure 4A compares the performance of INSIGHT, elnet, and XGBoost. DC was excluded from this analysis as it was not designed for combination predictions. INSIGHT demonstrated a significant performance improvement over existing methods. Detailed comparisons between INSIGHT and XGBoost are presented in Figure 4B, showing that while the methods were comparable to some drug response predictions, there was a subset of drug combinations where INSIGHT outperformed XGBoost.

**Figure 4.**
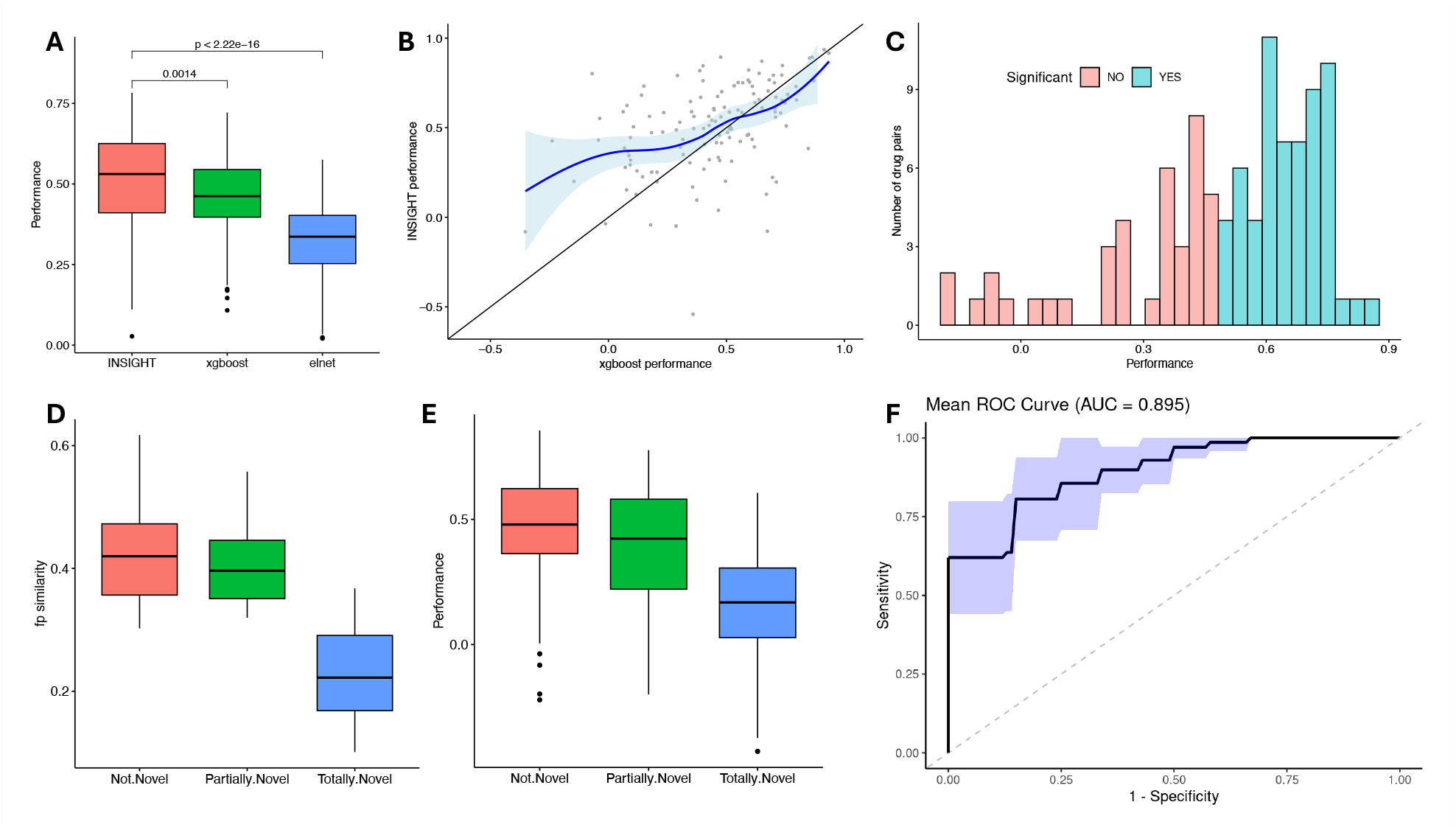
Performance on predicting drug sensitivity on DrugComb datasets. (A) Performance of predicting sensitivity for independent samples. (B) Detailed comparison of performance between INSIGHT and XGBoost. (C) Distribution of the performance of INSIGHT prediction on new drug pairs. A hypothesis test is done to label whether the correlation is significant. FDR < 0.1 is used for significance. (D) Performance of INSIGHT for drug pairs from three different similarity categories. (E) Similarity between testing drug pairs and training drug pairs. (F) ROC evaluation of predicting INSIGHT performance based on the variance of sensitivity prediction.

We further evaluated INSIGHT’s performance in predicting the sensitivity of novel drug combinations. For the reasons discussed in the single-agent modeling section, elnet and XGBoost were not tested. Out of all drug combinations with over 50 cell lines, 100 were randomly selected for independent testing. Figure 4C shows INSIGHT’s performance assessed by the Pearson’s correlation. Our analysis showed that 84 drugs exceeded a performance cutoff of 0.3 on the correlation coefficients and a subset of 64 also passed a statistical significance at the FDR cutoff of 0.1. We note that the limited number of cell lines tested in the DrugComb dataset makes it challenging to reach statistical significance for some of the drug combinations.

We also assessed the relationship between prediction performance and drug structure similarity, categorizing drug combinations as not novel, partially novel, and totally novel drugs. Not novel drugs involved novel combinations where both drugs were in the training set; partially novel pairs included drug pairs where only one drug in the combination was in the training, and totally novel pairs involve entirely new drug combinations with no drugs in the training set. Figure 4D depicts INSIGHT’s performance across these categories, while Figure 4E shows the similarity measures of the drug fingerprints. Drug combination similarity was defined by the average 2D Tanimoto index between the four single drug comparisons (see Methods) between the tested drug combination and the drug pairs in the training dataset. As expected, drug pairs with the highest similarity to the training data performed the best and totally novel pairs were the most challenging to predict. Not novel and partially novel drug pairs achieved similar performance because of the comparable drug structure similarity. For not novel pairs, we found a strong correlation between the performance and the variance of the predicted sensitivity (Supplemental Figure S8). This allowed us to use the variance to assess the performance of the predictions by applying a 10-fold cross validation using a linear regression model (using the rlm function in the MASS package, version 7.3-55). Figure 4F shows the ROC curve characterized by a high predictability of performance with an AUC of 0.895.

We note that the relationship between variance and performance remains valid for partially novel drug pairs, but not for totally novel pairs (Supplemental Figure S8). Indeed, for partially novel drug pairs a similar predictive performance analysis indicated a high predictability with an AUC of 0.892, as shown in Supplemental Figure S9.

### INSIGHT reveals pathway activity in response to drug treatments

In our model, the embedded pathways enabled us to evaluate the different pathways’ impact on drug response. We prioritize pathways by evaluating the correlation between pathway activities and predicted drug sensitivity across various cancer cell lines. As shown in Supplemental Figure S10, this correlation generally correlates positively with pathway size, indicating that larger pathways, encoding more general biological functions, tend to have stronger correlation with drug response. To overcome this bias, we considered the hierarchical structure of pathway relationships and adopted a method similar to the DrugCell’s RLIPP score, which calculates a relative score between the pathway and all its connected sub-pathways. This scoring helps to prioritize pathways where the correlation between pathway activity and drug sensitivity is not primarily driven by the pathway itself, but by some of its sub-pathways.

We first examined whether the pathways identified by INSIGHT are related to the MOA of drugs by checking if the pathways had the target genes included. Since drugs often have complex MOAs and typically present with off-target effects, we calculated the correlation between drug sensitivity and gene essentiality data as measured by the CRISPR screens from DepMap [26]. We filtered based on the correlation between drug sensitivity and the essentiality of target genes to identify drugs with well-defined targets in a data-driven way. We assumed that a high positive correlation between drug sensitivity and target gene essentiality increases our confidence that the annotated target gene was the main target of the drug. Of all the PRISM drugs 1,166 had annotated targets. After applying a correlation coefficient cutoff of 0.2 and a p-value of 0.01, we found 105 drugs with significant correlation between drug sensitivity and the target gene’s essentiality. For drugs with multiple annotated targets, the cutoff was applied to the target with the highest correlation, based on the rationale that this approach would help identify the drug’s primary target. For each of these 105 drugs, we identified the pathways using the INSIGHT model with correlation score and relative score cutoffs of 0.1 and 1, respectively. Of the 105 drugs, 90 (85.7%) have INSIGHT pathways containing the target gene from the Reactome pathway annotation, thus confirming that the model was able to identify the pathways related to the MOA of the drugs. We note that it is important to highlight that the model included only the drug’s structure and sensitivity profile, without incorporating any prior information about the drug’s target.

Next, we examined if we could classify drugs based on the pathway activity profiles. We analyzed the relative scores of all drugs and generated a low-dimensional reduction using UMAP to represent the relationships between compounds (see Methods for details). We hypothesized that drugs in proximity would share similar MOA and highlighted several instances from the PRISM screen where multiple drugs targeted the same pathways. Figure 5A shows the UMAP plot of EGFR inhibitors from the PRISM drugs. Indeed, we found that EGFR inhibitors that were FDA-approved or in Phase 2 or later clinical trials were all clustered together and tended to be more extensively studied, as evidenced by the higher number of articles available in PubMed. In contrast, most of the drugs annotated as EGFR inhibitors which were scattered in the plot were investigational compounds or in early-stage (phase 1) trials. These drugs may have off-target effects and their MOA might not be completely understood and thus being mislabeled as EGFR inhibitors. This could explain why they do not cluster with the well-studied EGFR inhibitors.

**Figure 5.**
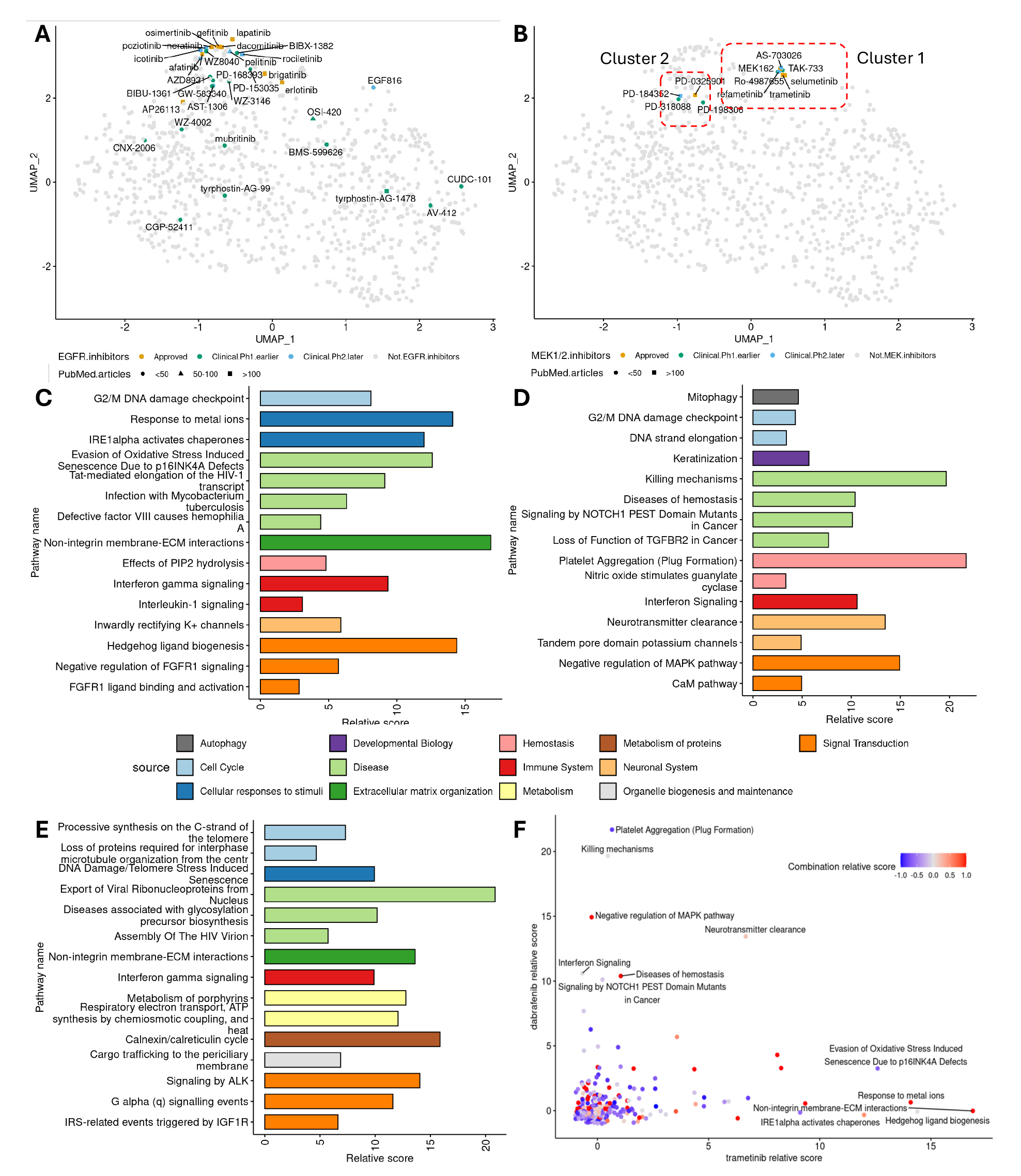
MOA interpretation of drugs. (A) UMAP analysis of pathway relative score for EGFR inhibitors. (B) UMAP analysis for MEK inhibitors. (C) Top 15 pathways from trametinib. (D) Top 15 pathways from dabrafenib. (E) Top 15 pathways from trametinib and dabrafenib combination. (F) Scatter plot of pathway scores between trametinib and dabrafenib. Color is coded by the relative score of combination therapy.

In Figure 5B, we show a detailed map of MEK 1/2 inhibitors, revealing two main clusters. Cluster 1 is composed of small molecule MEK inhibitors, with the majority being FDA-approved compounds like trametinib, selumetinib, and MEK162, all of which have shown increased efficacy in the presence of BRAF mutations. The other cluster comprises of four MEK inhibitor compounds, one of them, PD-184352 is among the first-generation MEK inhibitors to reach clinical trials [27]. The other three compounds, PD-0325901, PD-318088, and PD-198306 are structural analogs of PD-184352 [27–29] and it makes sense that they are in proximity in the UMAP plot. Interestingly there were other compounds in proximity of both EGFR and the MEK inhibitor clusters suggesting that they may have related MOAs and potential indications.

Next, to examine the model’s ability to reveal relevant pathways to drug response, we considered the combination therapy of trametinib and dabrafenib approved for several indications, including BRAF mutated metastatic melanoma and non-small cell lung cancer. We considered these two drugs because in addition to being well-studied regarding their MOA and adverse effects, they were also included as single agents in the PRISM screen. We built models and prioritized pathways to predict response to both single and combination therapy of trametinib and dabrafenib.

First, we examined the model built for trametinib, a small molecule inhibitor of MEK1 and MEK2. Pathways were prioritized using cutoffs of 0.1 on absolute pathway correlation scores and 1 on relative scores, identifying 41 pathways (Supplemental Table S1). Figure 5A shows the top 15 pathways, with non-integrin membrane-ECM interactions as one of the top pathways associated with drug response. There are several studies demonstrating trametinib’s influence on ECM function. Trametinib has been linked to decreased CD44 expression in mesothelioma cells, affecting ECM signaling pathways that encourage tumor growth [30]. Trametinib combined with dabrafenib has been shown to reduce ECM invasion in melanoma cells by downregulating PTTG1 [31], pathways also highlighted in our combination treatment models (Figure 5C). Furthermore, combining trametinib with afatinib improves treatment efficacy targeting ECM interactions [32]. Hedgehog ligand biogenesis also prioritized by our model, are areas of study for MEK inhibitors due to their role in tumorgenesis [33]. Additionally, it has been shown that trametinib may modulate interferon signaling to counter resistance and enhance therapy [34], a pathway identified by our model. Interestingly, there were several top pathways related to DNA damage that were related to drug response in our model. There is an interest in combining trametinib with DNA-damaging agents, the rationale being that inhibiting the MAPK/ERK pathway with trametinib could enhance the effectiveness of these treatments by preventing cancer cells from effectively repairing DNA damage. For example, combining trametinib with TAK-981, induces DNA damage accumulation in KRAS-mutant cancers, leading to tumor regression and apoptosis through inhibition of DNA repair proteins such as Rad51 and BRCA1 in preclinical models [35]. Our model supports the rationale of combining trametinib with DNA-damaging agents for a better therapeutic benefit. Interestingly, inwardly rectifying potassium (K+) channels, identified as one of the top pathways, plays role in affecting the contractile function of lymphatic vessels and may lead to lymphedema, which is a documented adverse event in patients taking trametinib, either as a standalone therapy or in combination with dabrafenib [36].

Next, we evaluated the pathway activity to response to dabrafenib applying the same cutoffs to the pathway scores and found several pathways related to the MOA including pathways directly linked to MEK targets, such as negative regulation of MAPK pathway and RAS mutant signaling (Supplemental Table S2). In addition, similar to the trametinib model we identified pathways related to interferon signaling and DNA damage. Keratinization, a top pathway prioritized by our model may be related to hyperkeratosis, a documented side effect of Dabrafenib’s [37]. Bleeding, another side effect of Dabrafenib, may be related to the platelet aggregation and hemostasis diseases pathways identified by our model [38].

We further examined the pathways of the trametinib and dabrafenib combination using the model trained on DrugComb data (Figure 5C). We identified 33 pathways with the same cutoffs, with 5 overlapping with those of trametinib or dabrafenib (Supplemental Table S3). Non-integrin membrane-ECM interactions appeared in both the trametinib, and combination model as documented previously [30]. Not surprisingly, as both interferon gamma signaling and DNA damage pathways were prioritized for both single treatments, they were also selected in the combination model. Interestingly, ALK signaling pathway was uniquely identified for the combination treatment, although there is no established evidence of direct effect of trametinib and dabrafenib on the ALK pathway. However, the inhibition of the MAPK/ERK pathway could potentially influence other pathways such as the ALK pathway by signaling crosstalk as suggested by our model. The combination therapy of dabrafenib and trametinib changes the adverse event profile, making certain single agent side effects such as hyperkeratosis less frequent [39]. This change in the adverse event profile may be reflected in the lack of some pathways related to single agent side effects in our combination therapy model.

Furthermore, we explored the relative scores of single agent treatments compared to the combination effect of the drugs by plotting the trametinib and dabrafenib pathway scores with color code for the combination therapy effects (Figure 5E). The unique pathway activity score profiles of the drugs in our models suggests that the enhanced drug efficacy was accomplished mostly by the complementary pathway inhibition effect of the combination therapy. The fact that the single agent treatments are targeting the different components of the interferon signaling pathways in our model (Figure 5E) may suggest an enhanced effect on immune cell modulation for the combination treatment therapy [40].

Next, to evaluate our ability to distinguish between the different combination sensitivity profiles we applied the INSIGHT models to make predictions to all combinations available from DrugComb across all cancer cell lines. We also included all possible novel combinations of the top 50 drugs that were most frequently studied by DrugComb. Based on a UMAP analysis we found very different predicted sensitivity profiles for the combinations of a drug with different partners. For example, we considered dasatinib and sorafenib combinations, being two of the top drugs tested in most cell lines in DrugComb database. As expected, some of the combinations with both dasatinib and sorafenib clustered together and had high correlation with the single agent treatments, suggesting that these combinations do not alter significantly the single agent sensitivity profiles (Supplementary Figure S11). However, there were combinations with very different sensitivity profiles from the single compound treatments. We highlighted two combinations, the dasatinib and sorafinib, and the dasatenib and lapatinib, as examples with very different profiles from the single agent treatments, both combinations having preclinical evidence for synergistic effects [41,42]. These examples illustrate how our predictions could be used to prioritize combination therapies based on their predicted sensitivity profiles for further experimental validation to evaluate potential synergistic effects.

## DISCUSSIONS

We developed an interpretable deep neural network to model drug sensitivity by combining multiomics data with structural drug information building on an earlier framework of DrugCell. Our model significantly extended the functionality of the earlier models, by generalizing the model for the integration of multiomics data and enabling predictions for combination therapies. The flexible framework of the model allows not only the integration of various omics data types, but also hierarchical functional classifications, such as pathway databases, gene signatures, gene ontologies, or other literature-based classifications to enable biological interpretability.

Our model is capable to perform in silico screening for both new cell lines and novel compounds accelerating the time needed to prioritize compounds for the different lineages in the preclinical drug discovery pipeline. The continuously expanding availability of molecular profiles of cell lines enables us to expand current drug screens by making predictions of drug response on the new samples based on their expression and mutation profiles for the cancer indication of interest.

The use of combination therapy in oncology is crucial to enhance treatment efficacy, overcome drug resistance and by targeting multiple pathways in cancer cells to reduce the likelihood of tumor adaptation [43–45]. Our method enables us to prioritize the most promising therapies given the large number of potential combinations of approved and investigational drug candidates. We demonstrated superior prediction accuracy compared to conventional state-of-the-art machine learning techniques and other biologically informed deep learning methods. We developed a classifier, based on the structure similarity of a novel compounds to the drugs in the training set, to assess our confidence to build well performing models. This allows us to increase the accuracy of our methodology by focusing only on novel compounds with some level of representation of similar drugs in the training set and to differentiate novel compounds suitable for in silico screenings from the compounds which need experimental testing for a more accurate assessment. Besides the predictions to identify the most efficacious drug combinations, we showed an application of our pathway prioritization methodology to enable MOA interpretation and to identify off target pathways by highlighting unexpected pathways that can be further examined for potential adverse effects.

One of the limitations of the current model is the relatively small number of cancer cell lines for the drug combinations in the DrugComb database. This limited sample size can lead to underfitting the deep learning model, potentially impacting its performance in predicting new combination drug response and the interpretation of pathway activity. However, as the predictions prioritize promising combinations and corresponding lineages, further drug screens can test these predictions, and the models iteratively retrained with new data to enhance accuracy and utility.

In recent work models built on large scale CRISPR screens in cancer cell lines were utilized to make on-target toxicity predictions using expression profiles in normal human tissue [46]. Similarly, in future work our deep learning model trained in cancer cell lines could predict the effect of both mono and drug combination therapy on normal cells using expression only to identify pathways with potential toxicity concerns.

As our model can make predictions for any novel compounds, it is possible to include the fingerprints of de novo compounds in combination with existing drugs in the model to maximize the efficacy and minimize adverse effects. Advances in deep learning approaches may enable us in the future to reverse engineer de novo compounds based on their molecular fingerprints [47] identified as efficacious by our model.

## METHODS

### PRISM data processing

PRISM released drug screening data in the second quarter of 2020 (20Q2), comprising both primary and secondary screening datasets [48]. The primary screening dataset involved subjecting various drugs to cell lines at fixed doses. Drug sensitivity was quantified based on the logarithmic fold change in the number of cells compared to control treatment.

In contrast, the secondary screening dataset encompassed a comprehensive range of drug doses and assessed sensitivity through the calculation of Area Under the Curve (AUC) values. AUC values provide a superior metric for characterizing drug sensitivity since they account for sensitivity variations across different doses. These values range from 0 to 1, with lower values indicating higher drug sensitivity. This dataset encompasses 1,452 drugs and 738 cell lines. For training and validating our deep learning models, we used the secondary screening data. We filtered out data points with AUC values greater than 1. For drugs screened in multiple batches, we retained only those with the highest number of cell lines tested. Specifically, we removed the drug doxycycline from the study, as it had the same screen ID in two batches. We also removed the drugs that are screened on less than 20 cell lines. Finally, the processed PRISM data consists of 447,287 data points with 1,397 drugs and 731 cell lines.

### DrugComb data processing

The raw data from DrugComb version 1.5 was obtained from https://drugcomb.fimm.fi/. We performed several filtering steps to prepare the data for our model. First, we removed drugs that lacked SMILES information. The SMILES information was collected from CHEMBL [49], PUBCHEM [50] and PRISM data. Next, we excluded cell lines without associated expression or mutation data from the DepMap dataset. For drug combination screens across multiple databases, we used the average sensitivity. The gene expression and mutation data were processed in the same manner as in the single agent model. Drug sensitivity values were uniformly scaled by a factor of 0.01, so they range from -1 to 1. We also removed the drug pairs that are screened on less than 10 cell lines. Finally the processed DrugComb data consists of 334,749 data points with 160 cell lines and 11,412 drug pairs from 280 drugs. For the drug combination fingerprints, we concatenated the fingerprints of the two drugs. Because the order of drugs does not impact actual sensitivity, each drug-cell line combination has two fingerprints, created by appending the drugs in different orders.

### Multiomics data processing

The raw gene expression and mutation data were obtained from the DepMap portal (21Q1 release, https://depmap.org/portal/). Gene expression data is measured as TPM on a log2 scale, while mutation data is summarized as a binary representation at the gene level. These data will be processed separately using the same procedure for single-agent and combo modeling due to different sets of cancer lines used. For the gene expression data, we used the FindVariableFeatures function from the Seurat package (version 4.4.0) to identify the top 8,000 genes with the most variable expression. In terms of mutation data, we retained only non-silent mutations with a frequency greater than 5%. Genes for both expression and mutation data were further filtered to include only those present in the Reactome pathways. Finally, 4,703 genes from the expression data and 5,202 genes from the mutation data were used for training the single-agent model. For the combo model, 4,334 genes from the expression data and 1,773 genes from the mutation data were utilized.

### Drug structure data processing

The SMILES annotations for drugs from the PRISM dataset were obtained via the DepMap portal. For drugs in the DrugComb database, SMILES data was sourced from PRISM, CHEMBL, and PubChem, resulting in over 3,000 drugs with available SMILES information. These SMILES were converted into fingerprints using the Infomax and Morgan fingerprint methods. The Infomax fingerprint, a continuous representation of 300 dimensions, is generated from a pre-trained Deep Graph Infomax model and is known for better sensitivity and synergy prediction performance, as described in [51]. The Morgan fingerprint was generated from SMILES to help assess the similarity between drugs. It was generated using RDKit python package (version 2023.09.2) with length 2,048 and a radius of 2. For drug combinations, the fingerprint is created by concatenating the fingerprints of drug pairs. As emphasized in the results section, each drug combination is represented in two ways by changing the order of concatenation since the order should not influence sensitivity outcomes.

### INSIGHT models

As illustrated in Figure 2A, we designed a two-level interpretable deep neural network to capture both sample-level and drug-level variance in drug sensitivity data. Each level shares the same architecture, as shown in Figure 2B, consisting of two components linked by an MLP network for prediction. For the interpretable neural networks, input data includes gene expression and mutations. These genes are linked through Reactome pathway networks. With the nodes in the top layer of the Reactome pathway network, a final pseudo layer was added to connect these top nodes. In this structure, each pathway is represented by 8 hidden neurons reflecting the pathway’s activity, which serve as inputs for the subsequent pathway level. The activity of pathway *i* can be represented as:

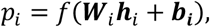

where ***W***_*i*_ is the weight vector, ***h***_*i*_ is the activity of hidden neurons, ***b***_*i*_ is the bias vector and *f* is the tanh activation function. For drug structure modeling, the drug fingerprint is embedded into an artificial neural network with three layers, each containing 128 neurons. The final output from this network is combined with the interpretable part and connected to an MLP with three fully connected layers of 128 neurons each for drug sensitivity prediction. Each hidden layer *l* can be represented by

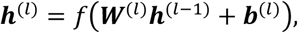

where ***h***^(*l*)^ is the hidden layer, ***W***^(*l*)^ the weight matrix, ***b***^(*l*)^ is the bias vector and *f* is the tanh activation function. The input layer ***h***^(()^ includes the embeddings from pathways and drug fingerprints.

The loss function incorporates a weighted sum of the root mean squared error (RMSE) of drug sensitivity and the activity fitting with the drug sensitivity, enhancing the interpretability of the drug’s MOA:

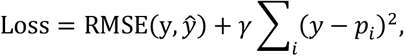

where y is the drug sensitivity data, ŷ is the predicted sensitivity, *p*_*i*_ is the activity of pathway *i* and *γ* is weighting parameter balancing the sensitivity prediction and pathway interpretability. Considering the scale of the two components, we selected *γ* as 0.001 to make the model have both predictability and interpretability. Parameter optimization was performed using the Adam optimization method [52] with a learning rate of 0.001. To mitigate overfitting and underfitting, 10% of the training data was reserved for validation. The number of training epochs was determined by monitoring performance on the validation samples.

### Similarity score calculation

The similarity score evaluates the structural similarity between a drug being tested and those used in the training set. This calculation involves two main steps. First, we identify the top 10 drugs from the training set that have the highest structural similarity to the drug in question, using the 2D Tanimoto index based on Morgan fingerprints. A minimum threshold of 0.15 is set for structural similarity to exclude drugs with distinctly different structures. Subsequently, we evaluated how well the drug under evaluation is represented by drugs with similar structures in the training set by the following equation.

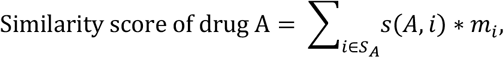

where *S*_*A*_ is the set of drugs which are similar to drug A in structure, *s*(*A, i*) is the Tanimoto similarity between drug A and *i*, and *m*_*i*_ is the model fitting performance of drug *i* which is calculated as the correlation coefficients between the predicting and fitting data. This score estimates the confidence in our model’s performance.

### Pathway score calculation

To assess pathway importance, we initially calculated a correlation score using Pearson’s correlation between the predicted sensitivity and the pathway hidden layer output. This score tends to favor higher-level pathways as they are closer to the output layer. Subsequently, we employed a method similar to the RLIPP score used in DC to evaluate subsystem importance. For each pathway, we constructed two linear models: one based on the hidden neurons of the pathway and another incorporating those of all its child pathways. The models’ response variable was the predicted sensitivity, and we determined the fitting correlation between the predicted sensitivity and the model output. The relative score is calculated as follows:

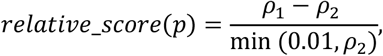

where *ρ*_1_ and *ρ*_2_ are the correlation coefficients of the first and second models, respectively. The cutoff of 0.01 in the denominator is used to prevent extreme values.

### UMAP analysis of pathway relative score

We evaluated the relative scores of all drugs and employed UMAP (R package v0.2.10.0) for a low-dimensional reduction to illustrate the relationships between compounds. From the PRISM dataset, we selected a subset of drugs accurately modeled by INSIGHT. We conducted a correlation analysis between the model-fitted sensitivity and the PRISM sensitivity. Using a fitting performance (Pearson correlation coefficient) cutoff of 0.2 and a p-value threshold of 1e-3, we identified 1,126 drugs for further analysis. In the EGFR inhibitors analysis, we restricted the number of annotated targets to a maximum of three to avoid overly complex inhibition interactions. For MEK1/2 inhibitors, we excluded drugs targeting genes other than MAP kinases. The drug labeled ‘MEK1-2-inhibitors’ (BRD-K12244279-001-05-8) was removed due to its generic name and our difficulty identifying the compound, while ‘nobiletin’ (BRD-K06753942-001-14-5) was removed because of lack of evidence about its target. Clinical information for the drugs was obtained from clinicaltrials.gov, and the number of PubMed articles was obtained by searching the drug names on https://pubmed.ncbi.nlm.nih.gov/.

### UMAP analysis of combo sensitivity prediction

To enhance our understanding of drug combination relationships, we used our model to expand the number of cell lines available for existing combinations, overcoming the sample size limitation in the current database. The raw data comprised 19,690 drug combinations with SMILES information before preprocessing. We identified the 50 drugs most frequently studied by DrugComb and created all possible combinations of these drugs as novel pairs. For each drug pair, we predicted the CSS across all cell lines of DrugComb and utilized the R package v0.2.10.0 to generate UMAP coordinates for all predictions. To examine the relationship between combinations involving dasatinib and dasatinib alone, we extracted the pathway activity for all related combinations from the model and compared it against the pathway activity of dasatinib using Pearson correlation. A lower correlation indicates that the combination’s MOA is more distinct from that of single-agent treatments. A similar analysis was conducted for sorafenib.

## ACKNOWLEDGEMENTS

We would like to express their sincere gratitude to Archana Iyer, Daniel Verduzco and Weilong Zhao from Genomics Research Center at AbbVie for their invaluable discussions and insightful suggestions for this research. We would also like to acknowledge the members of the Cancer Dependency Map Consortium for the use of the cancer cell line multiomics data, dependency and drug sensitivity data.

## CODE AVAILABILITY

The code of INSIGHT with demo data is available at https://github.com/xushiabbvie/INSIGHT.

## DATA AVAILABILITY

The processed data used for training is available at https://figshare.com/s/82a9d62c3da9f7176c91 and https://figshare.com/s/081d8df2a0c6143d2f45. It comprises processed PRISM data, DrugComb data, multiomics data, Reactome ontology networks and drug information.

## REFERENCES

[1] Hughes JP, Rees S, Kalindjian SB, Philpott KL. Principles of early drug discovery. Br J Pharmacol 2011;162:1239–49. 10.1111/j.1476-5381.2010.01127.x.

[2] Cook D, Brown D, Alexander R, March R, Morgan P, Satterthwaite G, et al. Lessons learned from the fate of AstraZeneca’s drug pipeline: a five-dimensional framework. Nat Rev Drug Discov 2014;13:419–31. 10.1038/nrd4309.

[3] Yang W, Soares J, Greninger P, Edelman EJ, Lightfoot H, Forbes S, et al. Genomics of Drug Sensitivity in Cancer (GDSC): a resource for therapeutic biomarker discovery in cancer cells. Nucleic Acids Res 2013;41:D955–961. 10.1093/nar/gks1111.

[4] Yu C, Mannan AM, Yvone GM, Ross KN, Zhang Y-L, Marton MA, et al. High-throughput identification of genotype-specific cancer vulnerabilities in mixtures of barcoded tumor cell lines. Nat Biotechnol 2016;34:419–23. 10.1038/nbt.3460.

[5] Zheng S, Aldahdooh J, Shadbahr T, Wang Y, Aldahdooh D, Bao J, et al. DrugComb update: a more comprehensive drug sensitivity data repository and analysis portal. Nucleic Acids Res 2021;49:W174–84. 10.1093/nar/gkab438.

[6] Ghandi M, Huang FW, Jané-Valbuena J, Kryukov GV, Lo CC, McDonald ER, et al. Next-generation characterization of the Cancer Cell Line Encyclopedia. Nature 2019;569:503–8. 10.1038/s41586-019-1186-3.

[7] Sharifi-Noghabi H, Zolotareva O, Collins CC, Ester M. MOLI: multi-omics late integration with deep neural networks for drug response prediction. Bioinforma Oxf Engl 2019;35:i501–9. 10.1093/bioinformatics/btz318.

[8] Baptista D, Ferreira PG, Rocha M. Deep learning for drug response prediction in cancer. Brief Bioinform 2021;22:360–79. 10.1093/bib/bbz171.

[9] Sapoval N, Aghazadeh A, Nute MG, Antunes DA, Balaji A, Baraniuk R, et al. Current progress and open challenges for applying deep learning across the biosciences. Nat Commun 2022;13:1728. 10.1038/s41467-022-29268-7.

[10] Tran KA, Kondrashova O, Bradley A, Williams ED, Pearson JV, Waddell N. Deep learning in cancer diagnosis, prognosis and treatment selection. Genome Med 2021;13:152. 10.1186/s13073-021-00968-x.

[11] Bhinder B, Gilvary C, Madhukar NS, Elemento O. Artificial Intelligence in Cancer Research and Precision Medicine. Cancer Discov 2021;11:900–15. 10.1158/2159-8290.CD-21-0090.

[12] Ching T, Himmelstein DS, Beaulieu-Jones BK, Kalinin AA, Do BT, Way GP, et al. Opportunities and obstacles for deep learning in biology and medicine. J R Soc Interface 2018;15:20170387. 10.1098/rsif.2017.0387.

[13] Wysocka M, Wysocki O, Zufferey M, Landers D, Freitas A. A systematic review of biologically-informed deep learning models for cancer: fundamental trends for encoding and interpreting oncology data. BMC Bioinformatics 2023;24:198. 10.1186/s12859-023-05262-8.

[14] Ma J, Yu MK, Fong S, Ono K, Sage E, Demchak B, et al. Using deep learning to model the hierarchical structure and function of a cell. Nat Methods 2018;15:290–8. 10.1038/nmeth.4627.

[15] Elmarakeby HA, Hwang J, Arafeh R, Crowdis J, Gang S, Liu D, et al. Biologically informed deep neural network for prostate cancer discovery. Nature 2021;598:348–52. 10.1038/s41586-021-03922-4.

[16] Doncevic D, Herrmann C. Biologically informed variational autoencoders allow predictive modeling of genetic and drug-induced perturbations. Bioinformatics 2023;39:btad387. 10.1093/bioinformatics/btad387.

[17] Kuenzi BM, Park J, Fong SH, Sanchez KS, Lee J, Kreisberg JF, et al. Predicting Drug Response and Synergy Using a Deep Learning Model of Human Cancer Cells. Cancer Cell 2020;38:672-684.e6. 10.1016/j.ccell.2020.09.014.

[18] Park S, Soh J, Lee H. Super.FELT: supervised feature extraction learning using triplet loss for drug response prediction with multi-omics data. BMC Bioinformatics 2021;22:269. 10.1186/s12859-021-04146-z.

[19] Milacic M, Beavers D, Conley P, Gong C, Gillespie M, Griss J, et al. The Reactome Pathway Knowledgebase 2024. Nucleic Acids Res 2024;52:D672–8. 10.1093/nar/gkad1025.

[20] Rogers D, Hahn M. Extended-Connectivity Fingerprints. J Chem Inf Model 2010;50:742–54. 10.1021/ci100050t.

[21] Holbeck SL, Camalier R, Crowell JA, Govindharajulu JP, Hollingshead M, Anderson LW, et al. The National Cancer Institute ALMANAC: A Comprehensive Screening Resource for the Detection of Anticancer Drug Pairs with Enhanced Therapeutic Activity. Cancer Res 2017;77:3564–76. 10.1158/0008-5472.CAN-17-0489.

[22] O’Neil J, Benita Y, Feldman I, Chenard M, Roberts B, Liu Y, et al. An Unbiased Oncology Compound Screen to Identify Novel Combination Strategies. Mol Cancer Ther 2016;15:1155–62. 10.1158/1535-7163.MCT-15-0843.

[23] Forcina GC, Conlon M, Wells A, Cao JY, Dixon SJ. Systematic Quantification of Population Cell Death Kinetics in Mammalian Cells. Cell Syst 2017;4:600-610.e6. 10.1016/j.cels.2017.05.002.

[24] Licciardello MP, Ringler A, Markt P, Klepsch F, Lardeau C-H, Sdelci S, et al. A combinatorial screen of the CLOUD uncovers a synergy targeting the androgen receptor. Nat Chem Biol 2017;13:771–8. 10.1038/nchembio.2382.

[25] Preuer K, Lewis RPI, Hochreiter S, Bender A, Bulusu KC, Klambauer G. DeepSynergy: predicting anti-cancer drug synergy with Deep Learning. Bioinformatics 2018;34:1538–46. 10.1093/bioinformatics/btx806.

[26] Tsherniak A, Vazquez F, Montgomery PG, Weir BA, Kryukov G, Cowley GS, et al. Defining a Cancer Dependency Map. Cell 2017;170:564-576.e16. 10.1016/j.cell.2017.06.010.

[27] Frémin C, Meloche S. From basic research to clinical development of MEK1/2 inhibitors for cancer therapy. J Hematol OncolJ Hematol Oncol 2010;3:8. 10.1186/1756-8722-3-8.

[28] Wu P-K, Park J-I. MEK1/2 Inhibitors: Molecular Activity and Resistance Mechanisms. Semin Oncol 2015;42:849–62. 10.1053/j.seminoncol.2015.09.023.

[29] Ripple MO, Kim N, Springett R. Acute Mitochondrial Inhibition by Mitogen-activated Protein Kinase/Extracellular Signal-regulated Kinase Kinase (MEK) 1/2 Inhibitors Regulates Proliferation. J Biol Chem 2013;288:2933–40. 10.1074/jbc.M112.430082.

[30] Cho H, Matsumoto S, Fujita Y, Kuroda A, Menju T, Sonobe M, et al. Trametinib plus 4-Methylumbelliferone Exhibits Antitumor Effects by ERK Blockade and CD44 Downregulation and Affects PD-1 and PD-L1 in Malignant Pleural Mesothelioma. J Thorac Oncol 2017;12:477– 90. 10.1016/j.jtho.2016.10.023.

[31] Caporali S, Alvino E, Lacal PM, Ruffini F, Levati L, Bonmassar L, et al. Targeting the PTTG1 oncogene impairs proliferation and invasiveness of melanoma cells sensitive or with acquired resistance to the BRAF inhibitor dabrafenib. Oncotarget 2017;8:113472–93. 10.18632/oncotarget.23052.

[32] Shen X, Jin X, Fang S, Chen J. EFEMP2 upregulates PD-L1 expression via EGFR/ERK1/2/c-Jun signaling to promote the invasion of ovarian cancer cells. Cell Mol Biol Lett 2023;28:53. 10.1186/s11658-023-00471-8.

[33] Macdonald JB, Macdonald B, Golitz LE, LoRusso P, Sekulic A. Cutaneous adverse effects of targeted therapies: Part II: Inhibitors of intracellular molecular signaling pathways. J Am Acad Dermatol 2015;72:221–36; quiz 237–8. 10.1016/j.jaad.2014.07.033.

[34] Cheng K, Zhou Z, Chen Q, Chen Z, Cai Y, Cai H, et al. CDK4/6 inhibition sensitizes MEK inhibition by inhibiting cell cycle and proliferation in pancreatic ductal adenocarcinoma. Sci Rep 2024;14:8389. 10.1038/s41598-024-57417-z.

[35] Kotani H, Oshima H, Boucher JC, Yamano T, Sakaguchi H, Sato S, et al. Dual inhibition of SUMOylation and MEK conquers MYC-expressing KRAS-mutant cancers by accumulating DNA damage. J Biomed Sci 2024;31:68. 10.1186/s12929-024-01060-3.

[36] Denton JS, Delpire E. Special collection on inward rectifying K+ channels. Am J Physiol - Cell Physiol 2023;324:C603–5. 10.1152/ajpcell.00457.2022.

[37] Peng C, Jie-Xin L. The incidence and risk of cutaneous toxicities associated with dabrafenib in melanoma patients: a systematic review and meta-analysis. Eur J Hosp Pharm Sci Pract 2021;28:182–9. 10.1136/ejhpharm-2020-002347.

[38] Lee LM, Feun L, Tan Y. A case of intracranial hemorrhage caused by combined dabrafenib and trametinib therapy for metastatic melanoma. Am J Case Rep 2014;15:441–3. 10.12659/AJCR.890875.

[39] Spain L, Julve M, Larkin J. Combination dabrafenib and trametinib in the management of advanced melanoma with BRAFV600 mutations. Expert Opin Pharmacother 2016;17:1031–8. 10.1517/14656566.2016.1168805.

[40] Vella LJ, Andrews MC, Pasam A, Woods K, Behren A, Cebon JS. The kinase inhibitors dabrafenib and trametinib affect isolated immune cell populations. Oncoimmunology 2014;3:e946367. 10.4161/21624011.2014.946367.

[41] Normann LS, Haugen MH, Hongisto V, Aure MR, Leivonen S-K, Kristensen VN, et al. High-throughput screen in vitro identifies dasatinib as a candidate for combinatorial treatment with HER2-targeting drugs in breast cancer. PLOS ONE 2023;18:e0280507. 10.1371/journal.pone.0280507.

[42] Cheng C-C, Chao W-T, Shih J-H, Lai Y-S, Hsu Y-H, Liu Y-H. Sorafenib combined with dasatinib therapy inhibits cell viability, migration, and angiogenesis synergistically in hepatocellular carcinoma. Cancer Chemother Pharmacol 2021;88:143–53. 10.1007/s00280-021-04272-8.

[43] Webster RM. Combination therapies in oncology. Nat Rev Drug Discov 2016;15:81–2. 10.1038/nrd.2016.3.

[44] Mokhtari RB, Homayouni TS, Baluch N, Morgatskaya E, Kumar S, Das B, et al. Combination therapy in combating cancer. Oncotarget 2017;8:38022–43. 10.18632/oncotarget.16723.

[45] Palmer AC, Sorger PK. Combination Cancer Therapy Can Confer Benefit via Patient-to-Patient Variability without Drug Additivity or Synergy. Cell 2017;171:1678-1691.e13. 10.1016/j.cell.2017.11.009.

[46] Shi X, Gekas C, Verduzco D, Petiwala S, Jeffries C, Lu C, et al. Building a translational cancer dependency map for The Cancer Genome Atlas. Nat Cancer 2024;5:1176–94. 10.1038/s43018-024-00789-y.

[47] Le T, Winter R, Noé F, Clevert D-A. Neuraldecipher – reverse-engineering extended-connectivity fingerprints (ECFPs) to their molecular structures. Chem Sci 2020;11:10378–89. 10.1039/D0SC03115A.

[48] Corsello SM, Nagari RT, Spangler RD, Rossen J, Kocak M, Bryan JG, et al. Discovering the anti-cancer potential of non-oncology drugs by systematic viability profiling. Nat Cancer 2020;1:235– 48. 10.1038/s43018-019-0018-6.

[49] Mendez D, Gaulton A, Bento AP, Chambers J, De Veij M, Félix E, et al. ChEMBL: towards direct deposition of bioassay data. Nucleic Acids Res 2019;47:D930–40. 10.1093/nar/gky1075.

[50] Kim S, Chen J, Cheng T, Gindulyte A, He J, He S, et al. PubChem 2025 update. Nucleic Acids Res 2025;53:D1516–25. 10.1093/nar/gkae1059.

[51] Zagidullin B, Wang Z, Guan Y, Pitkänen E, Tang J. Comparative analysis of molecular fingerprints in prediction of drug combination effects. Brief Bioinform 2021;22:bbab291. 10.1093/bib/bbab291.

[52] Kingma DP, Ba J. Adam: A Method for Stochastic Optimization 2017. 10.48550/arXiv.1412.6980.

