## Supplemental Figures and Methods for "INSIGHT: In Silico Drug Screening Platform using Interpretable Deep Learning Network"

### Supplemental files

Figure S1. Predicted sensitivity from DC using PRISM training data. (A) The graph shows the correlation between the average predicted sensitivity from DC (y-axis) and actual PRISM sensitivity for each drug (x-axis). The strong correlation demonstrates DC's capability to distinguish differences among drugs. (B) This panel presents examples of predicted sensitivity by DC alongside PRISM sensitivity for three different drugs. While DC accurately identifies the relative sensitivity of the three drugs, the variation in sensitivity across various cancer cell lines (y-axis) is limited.

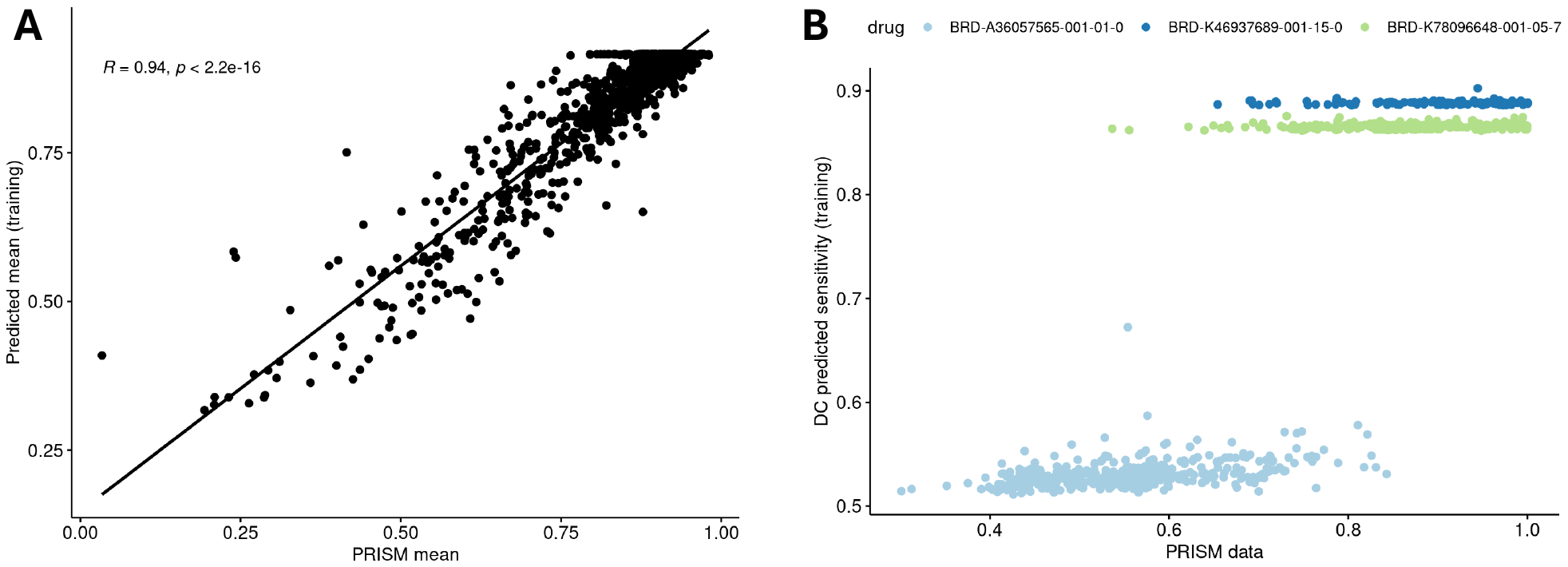

Figure S2. Feature selection between drug sensitivity and multiomics feature. (A) Coefficients of expression and mutation features from LASSO analysis predicting drug sensitivity. (B) The number of selected expression and mutation features from the LASSO analysis. (C) Percentage of drugs with at least one expression or mutation features selected by the LASSO analysis.

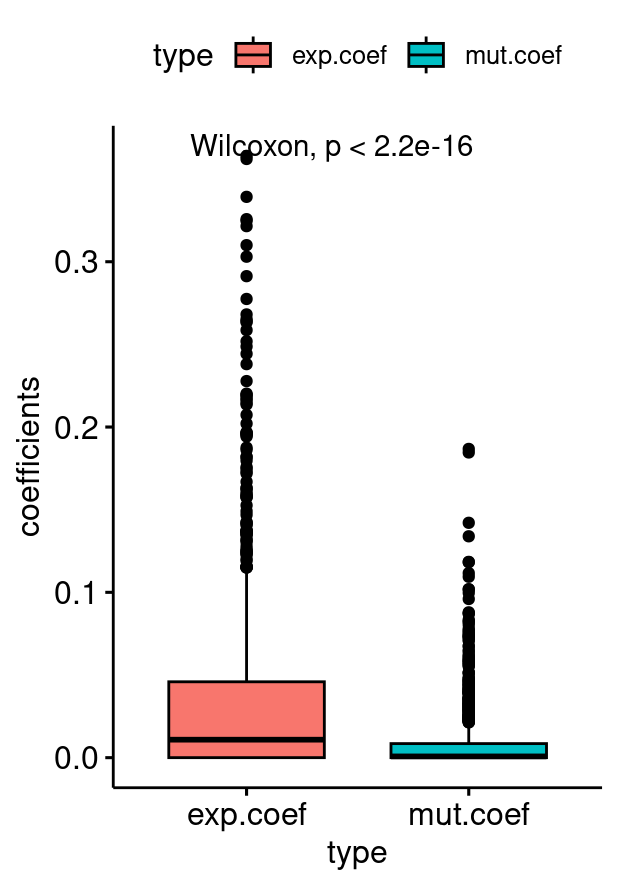

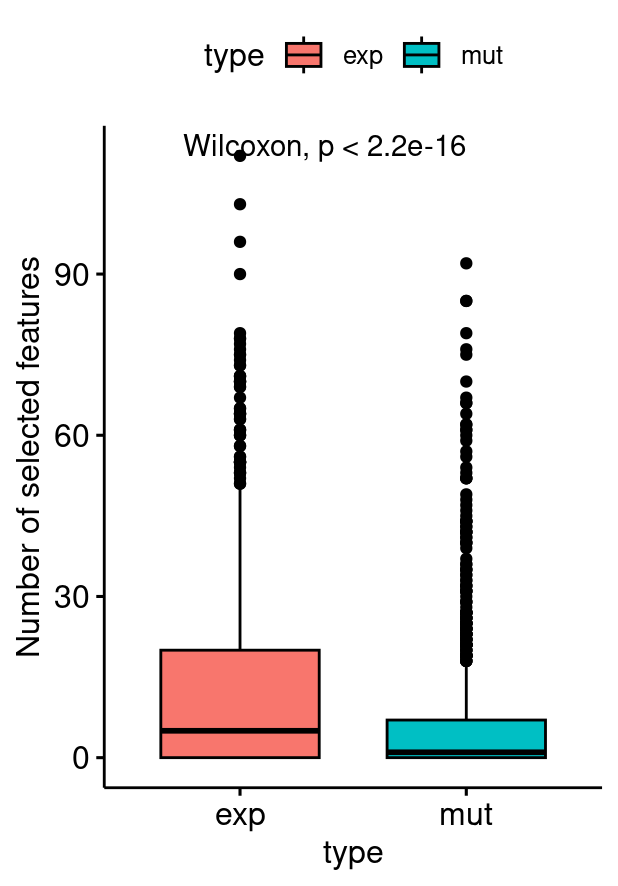

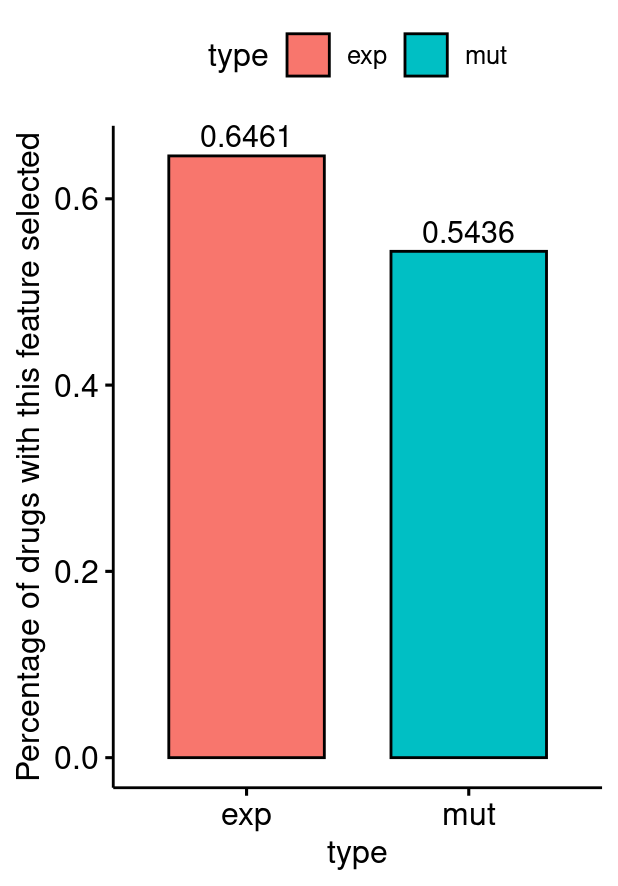

**B**

**C**

**A**

Figure S3. Pairwise correlation distribution of XGBoost.

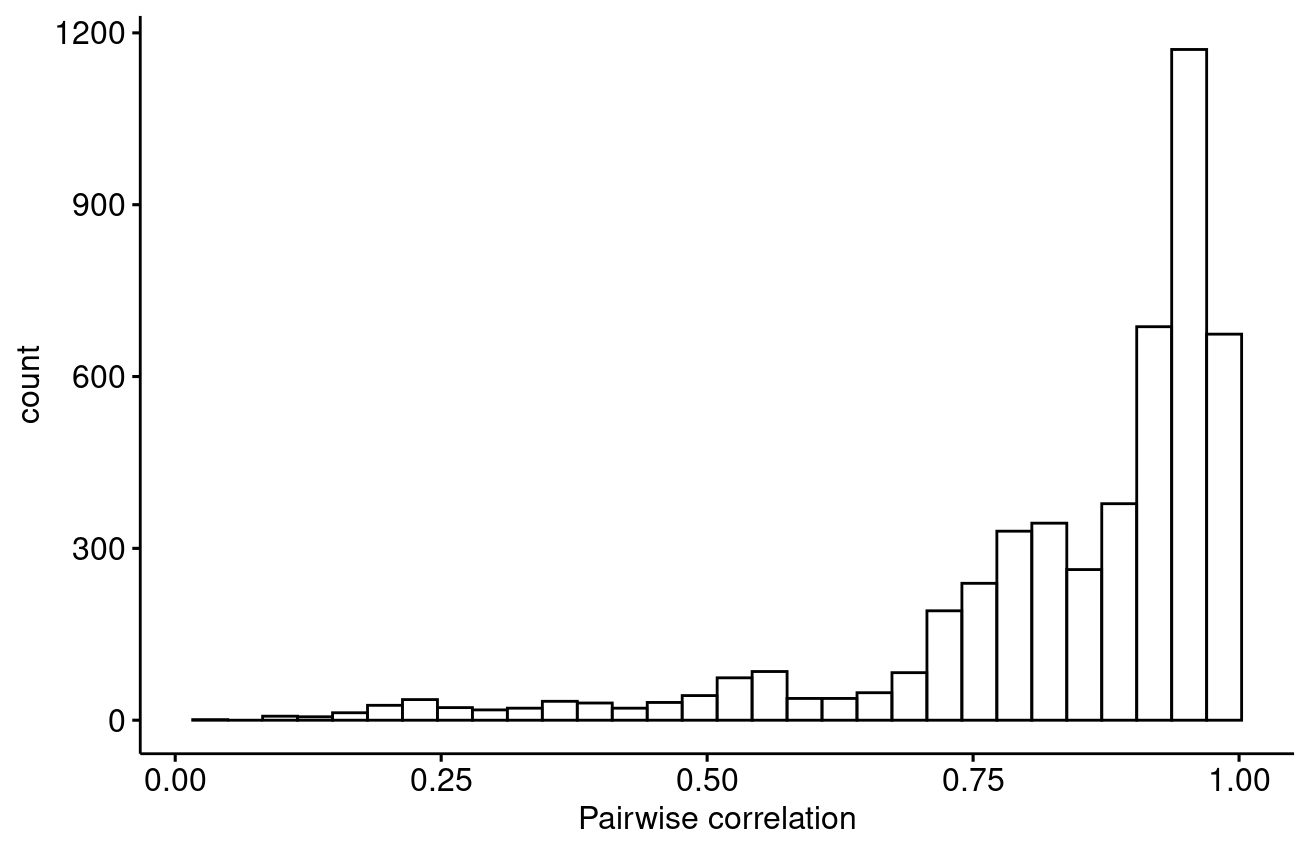

Figure S4. Pairwise correlation distribution of DC.

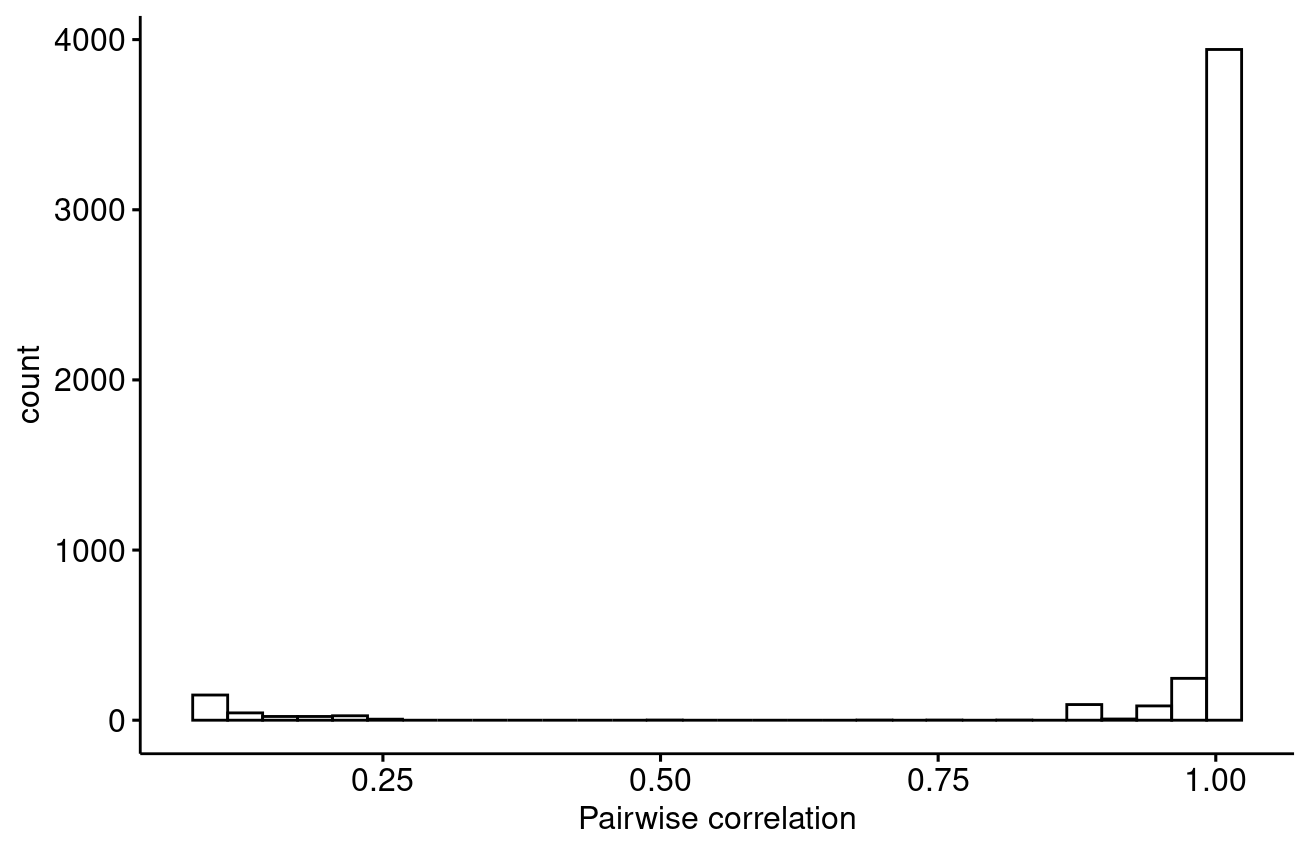

Figure S5. Correlation of BRD-A06627858-236-03-0 and BRD-A49035384-003-28-9 on (A) PRISM measured sensitivity, (B) INSIGHT predicted sensitivity, (C) XGBoost predicted sensitivity and (D) DC predicted sensitivity.

**B**

**A**

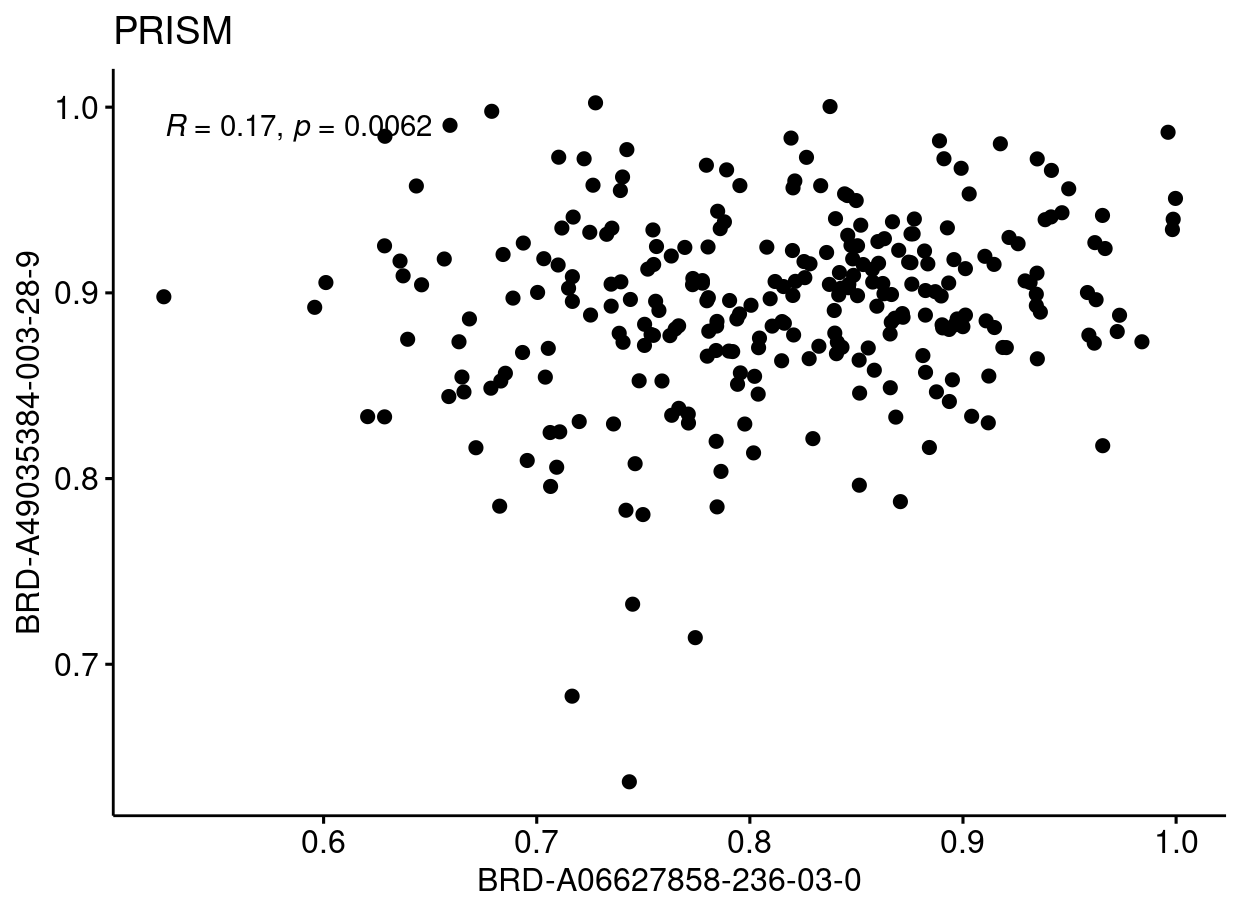

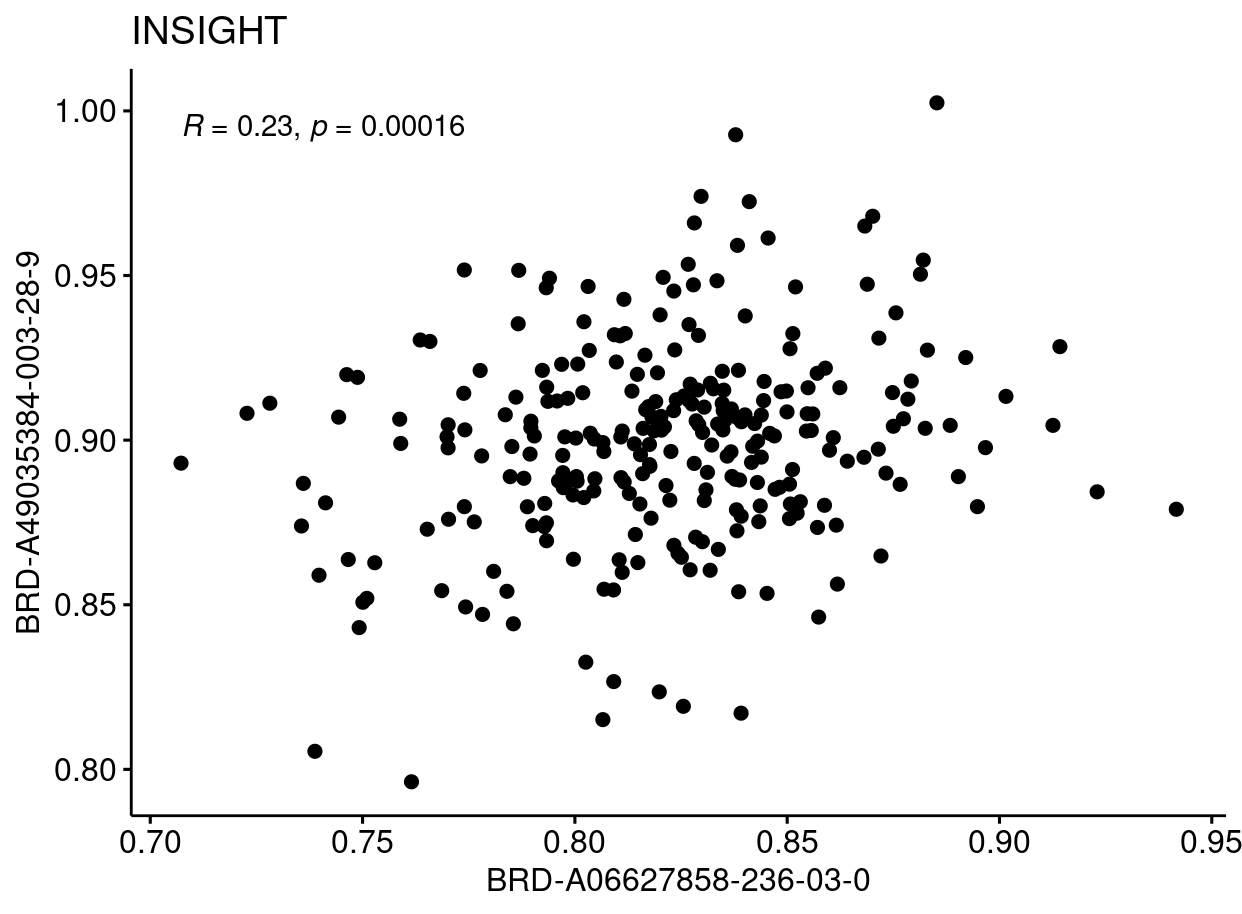

**C**

**D**

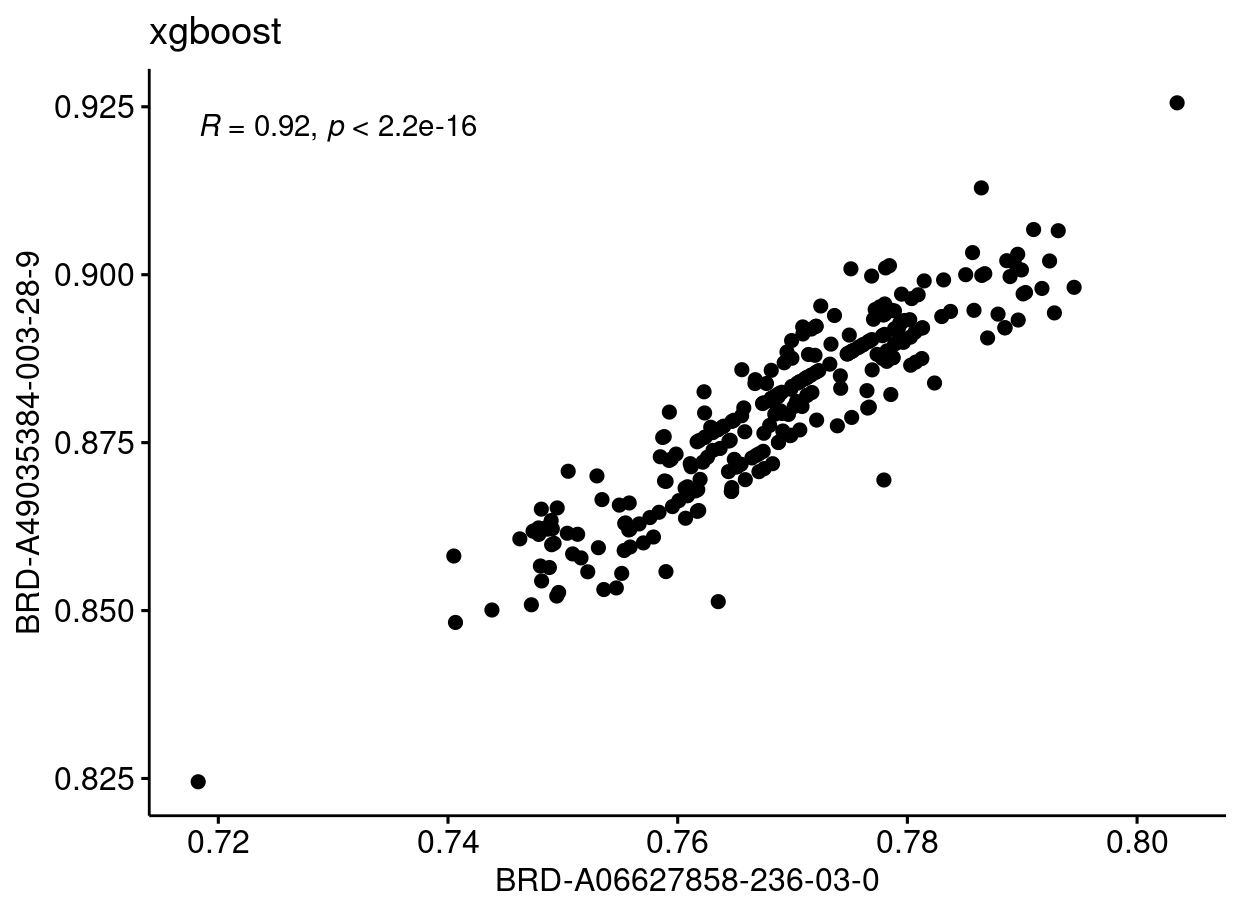

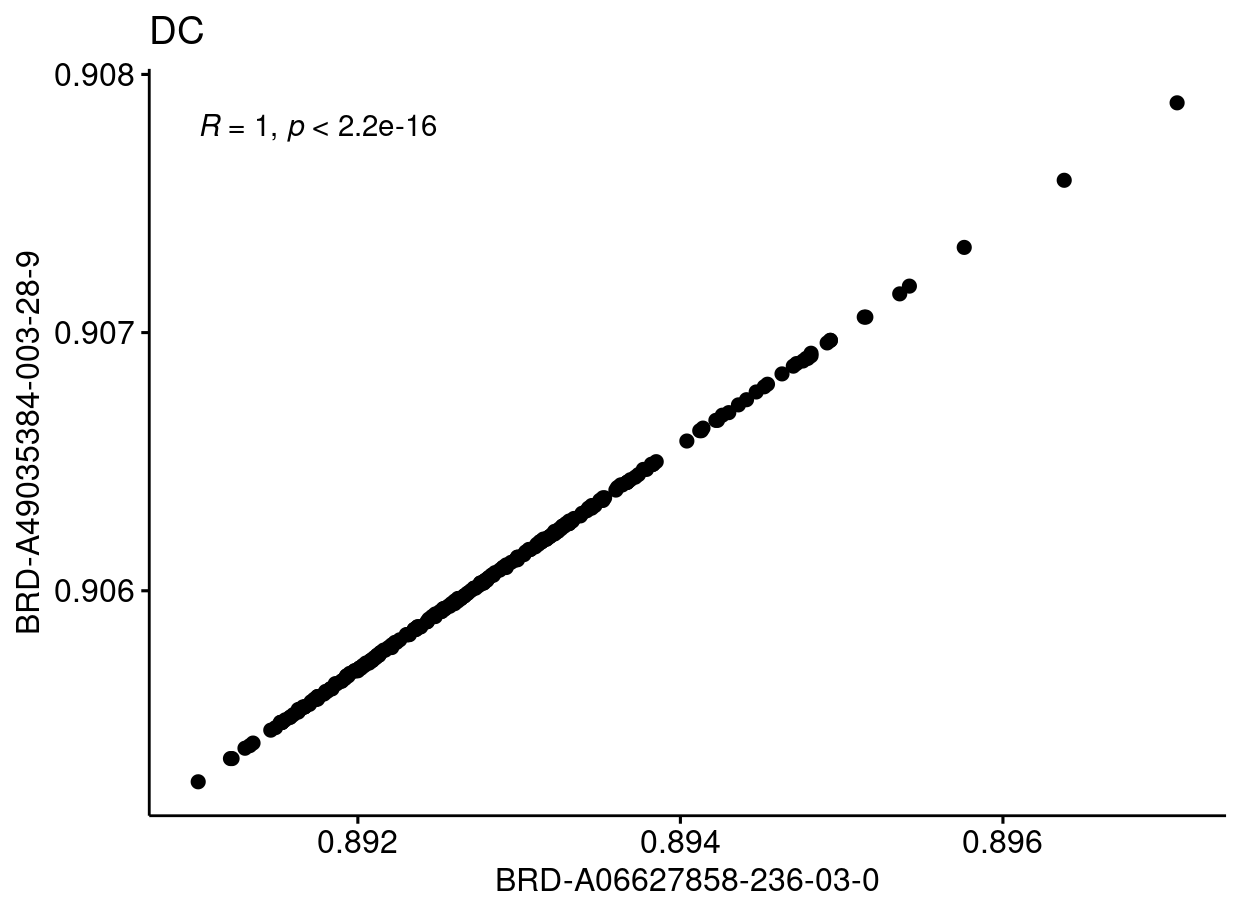

Figure S6. Example of vinblastine prediction

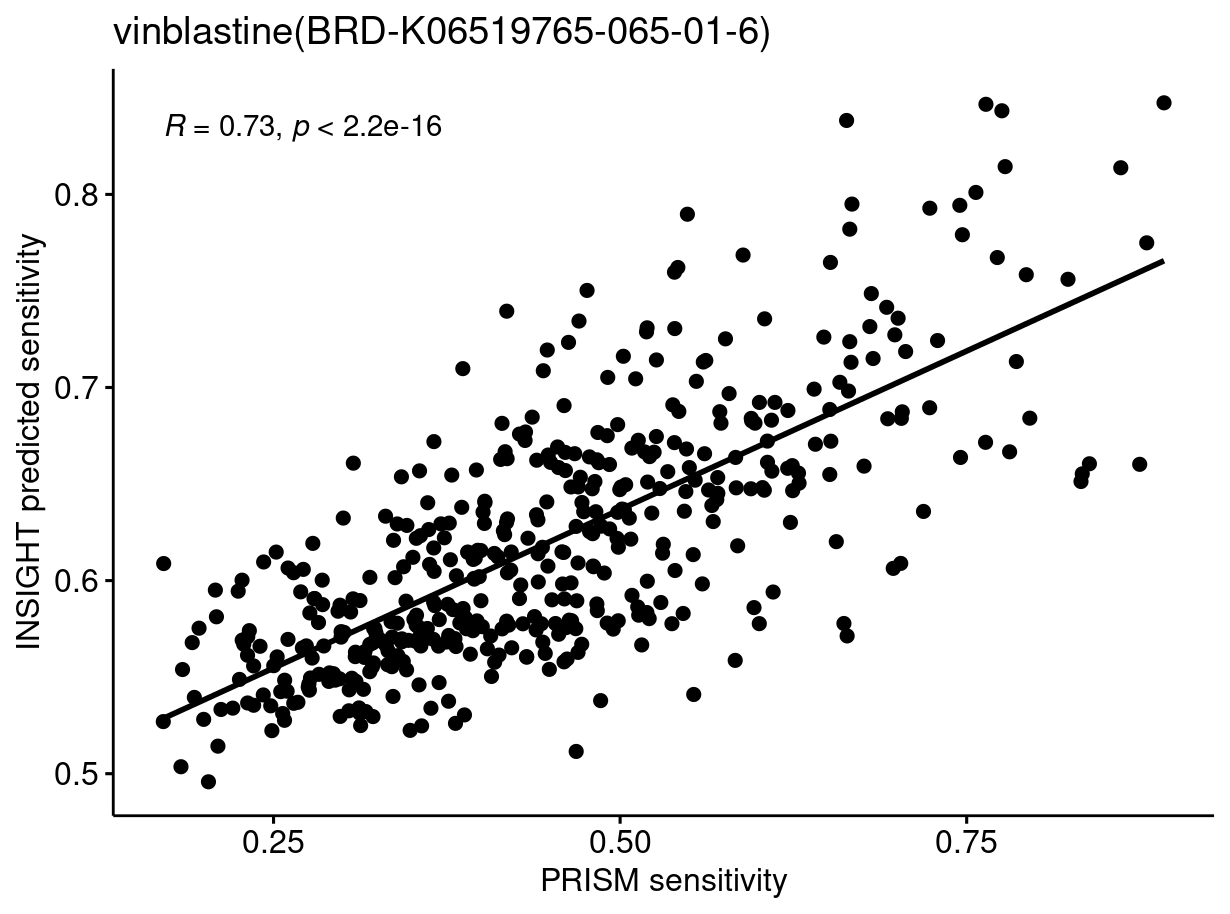

Figure S7. Correlation between INSIGHT prediction performance, similarity measure and variance of predicted sensitivity. (A) Morgan fingerprint similarity with higher values means more similar structure. (B) Infomax fingerprint distance with higher values means more different structure. (C) Correlation between variance of predicted sensitivity and INSIGHT performance.

**A**

**B**

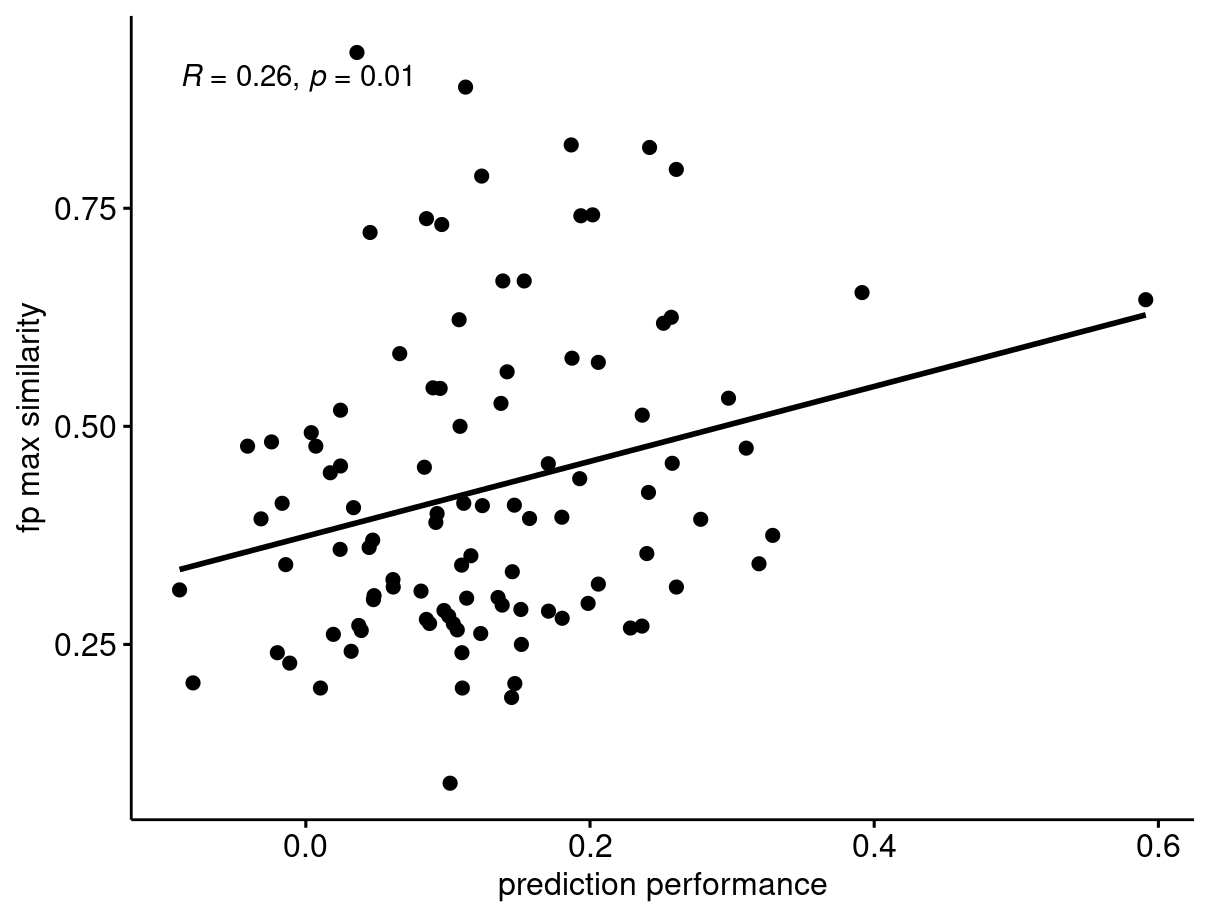

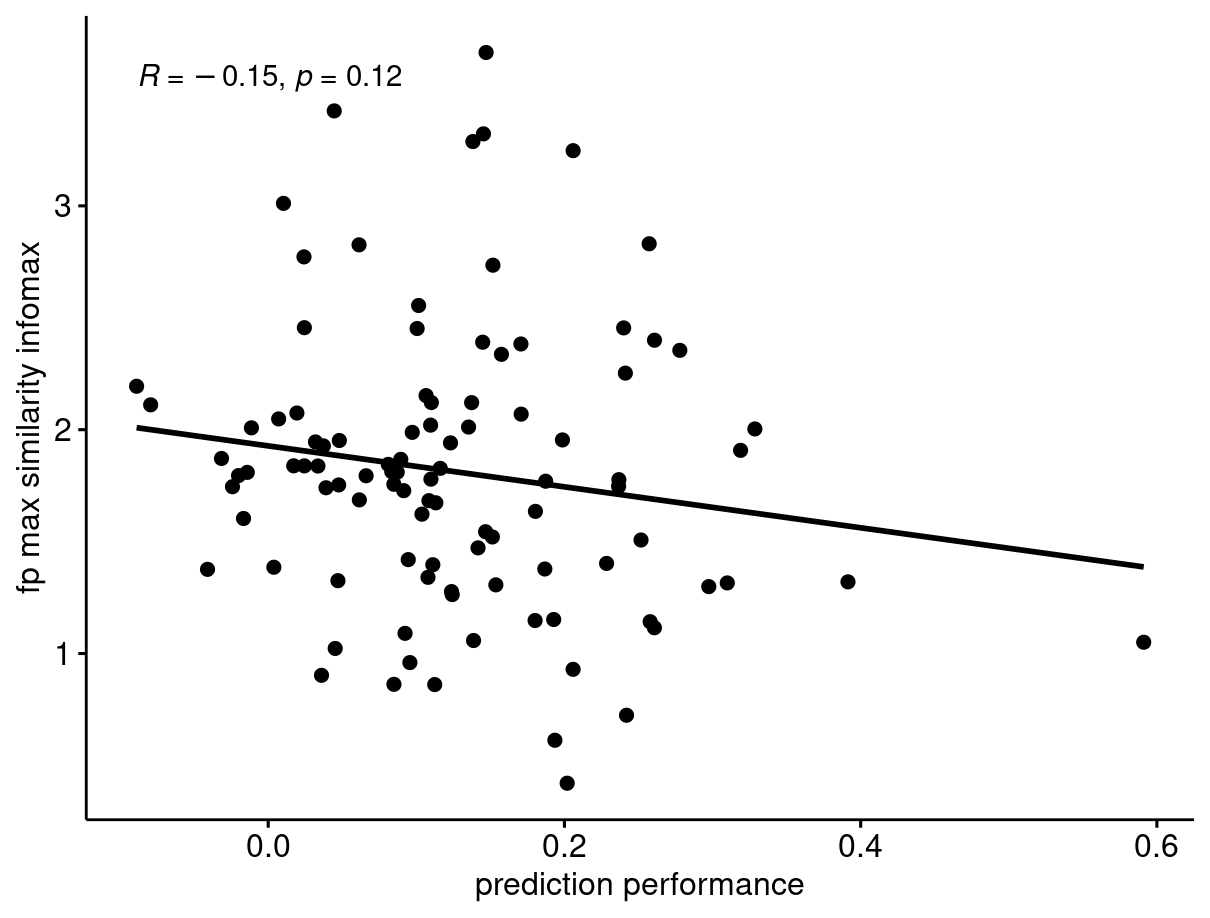

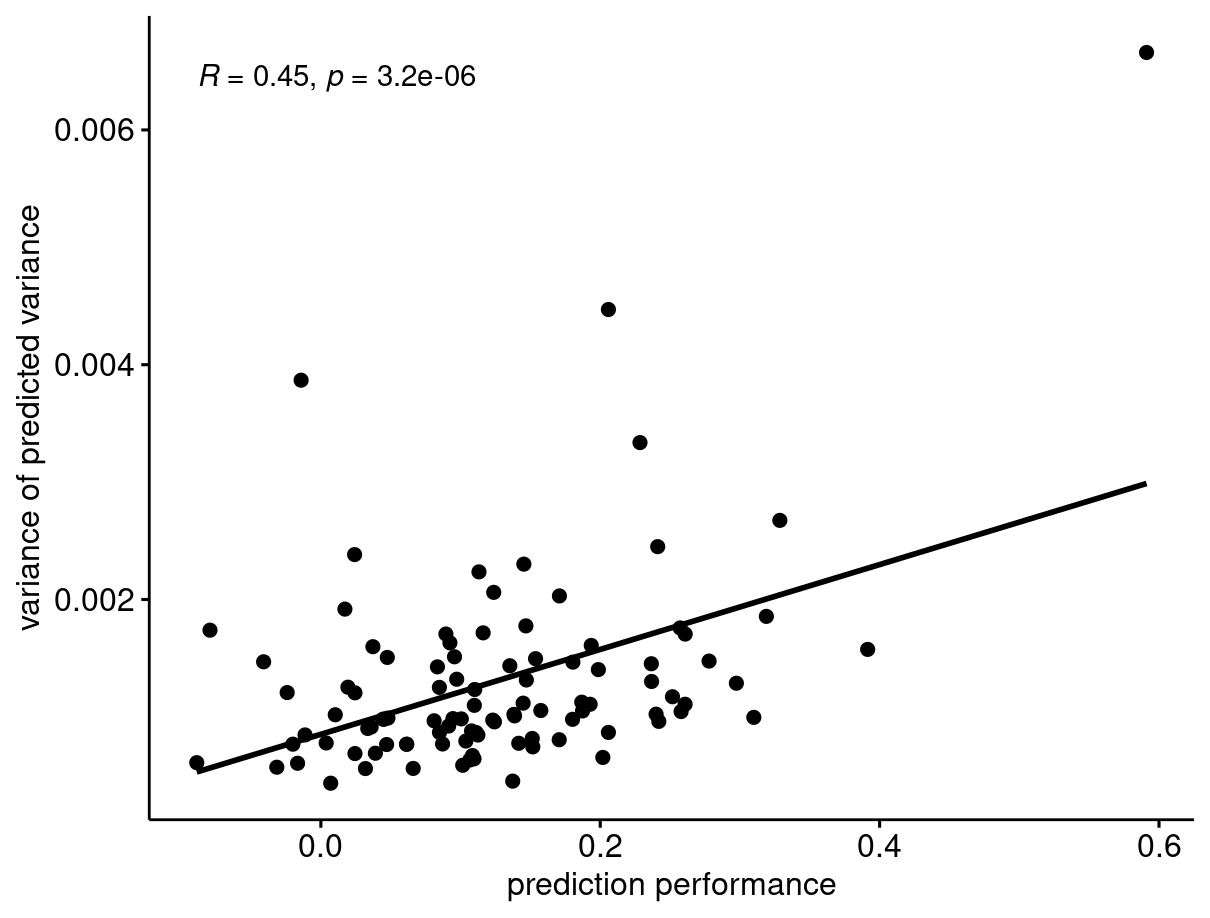

**C**

Figure S8. Relationship between the variance of predicted sensitivity and the performance for not novel (A), partially novel (B) and totally novel (C) drug pairs.

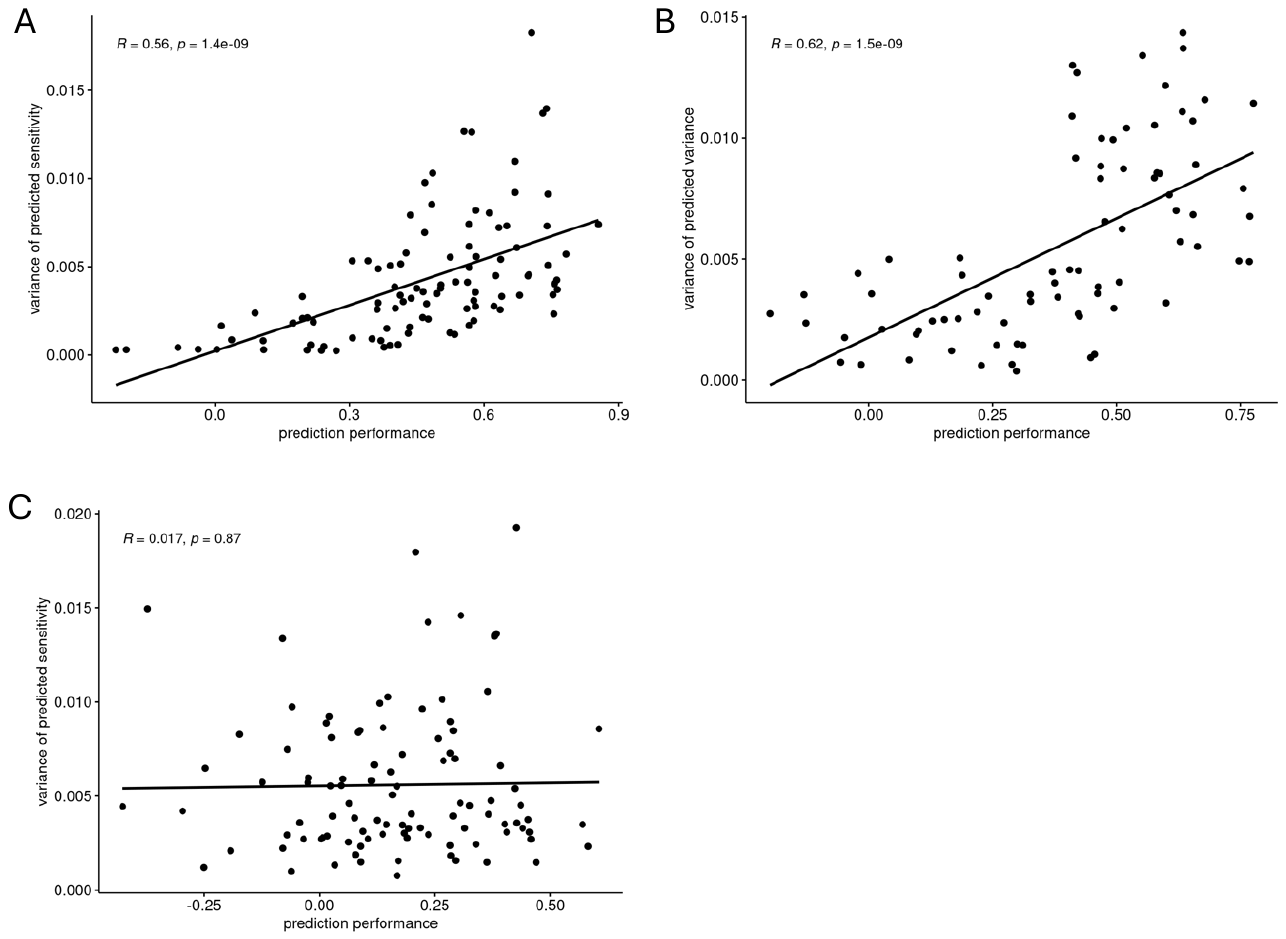

Figure S9. Performance of model to predict the performance of paritally novel drug pairs

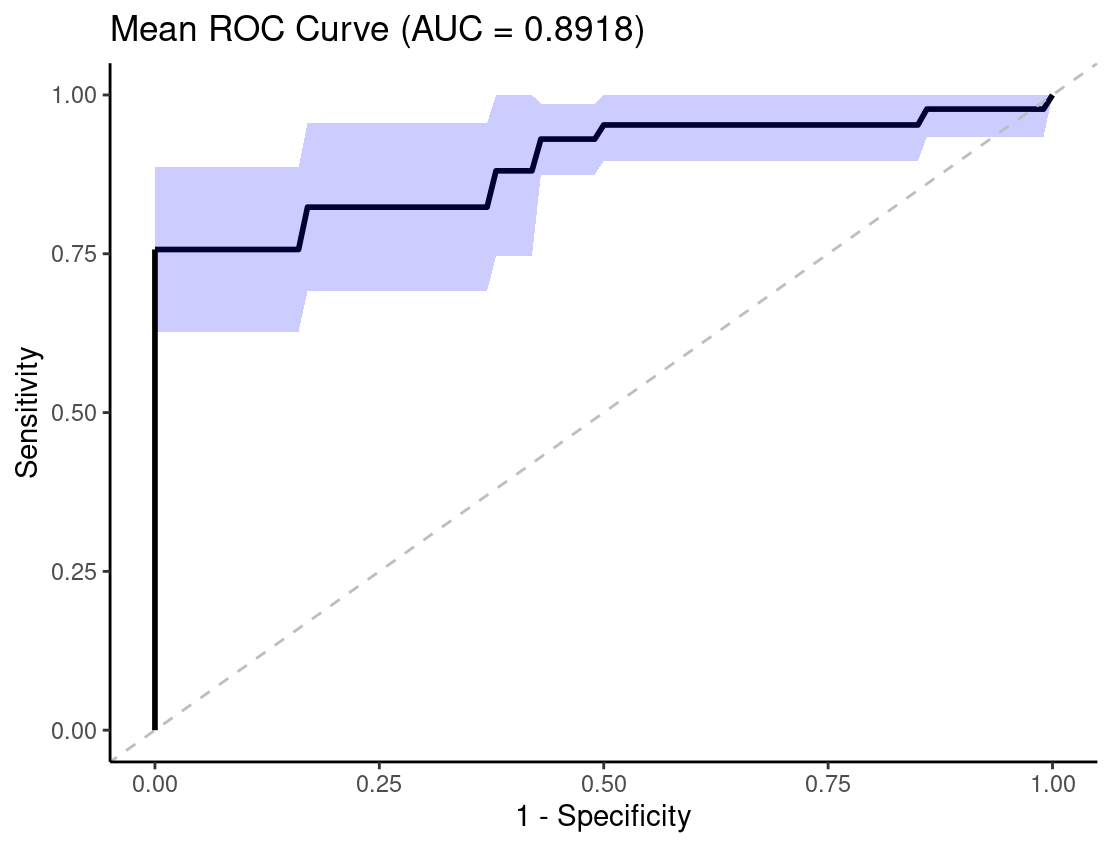

Figure S10. Correlation between pathway correaltion score and pathway size

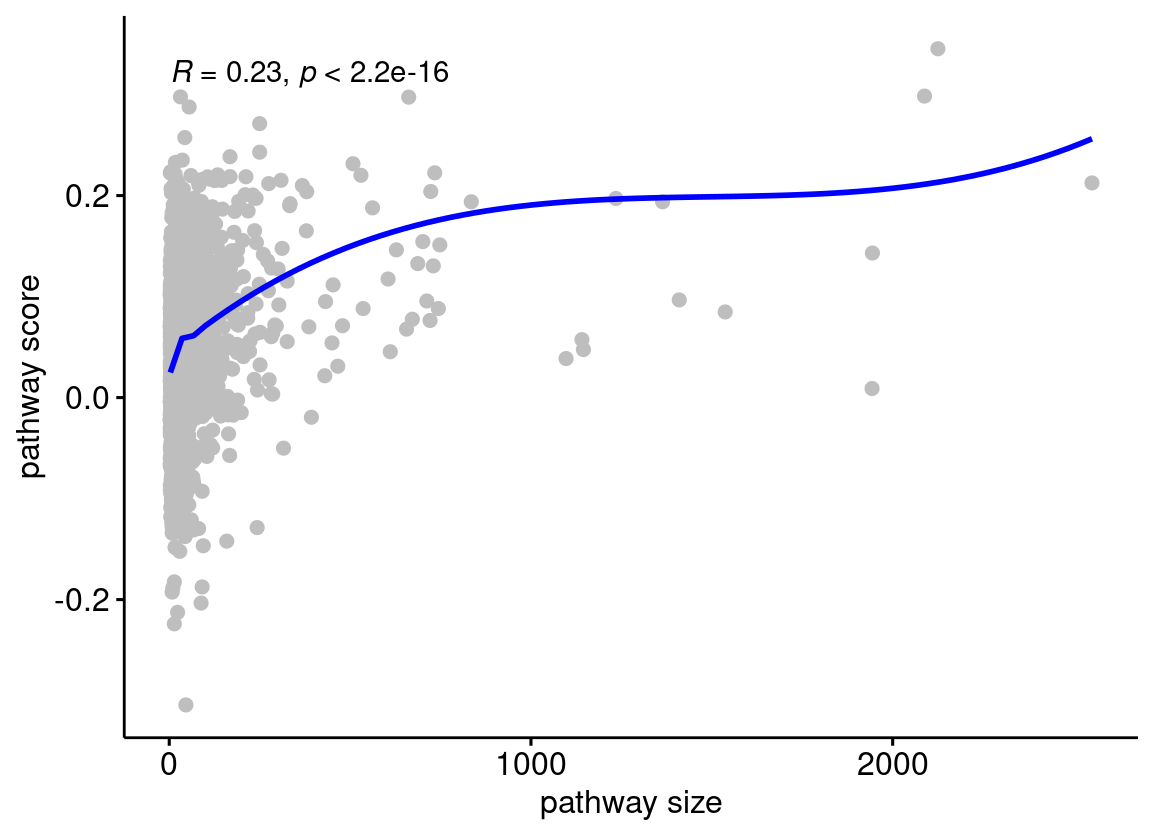

Figure S11. UMAP analysis of predicted drug senstivity of combinations.

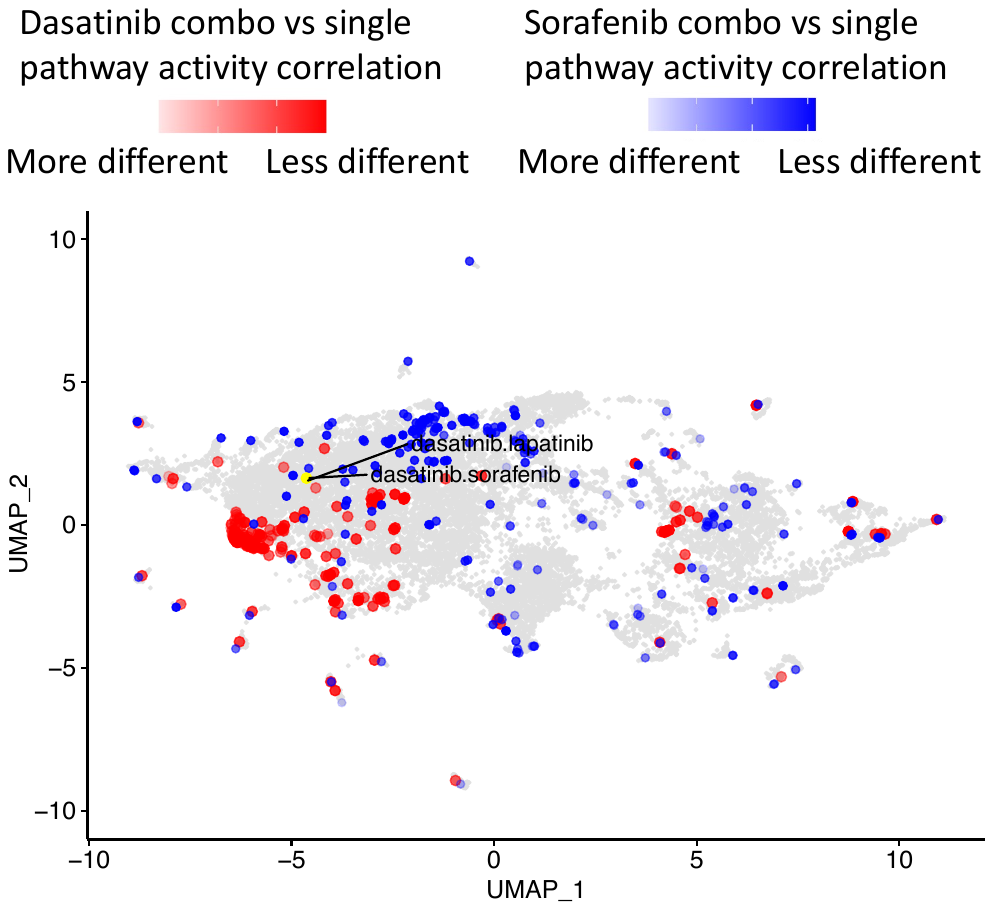

#### Supplemental Methods

##### S1. Prediction of drug sensitivity from elastic-net

We compared the performance of INSIGHT with elastic-net to predict drug sensitivity. The feature matrix consisted of combined gene expression, mutation, and drug fingerprint data. In the context of a linear elastic-net model, the sensitivity of drug A on cell line N can be expressed as:

$$y_{A,N}=\sum_{i} w_{o,i}x_{N,i}+\sum_{k} w_{p,k}f_{A,j}+c,$$

where $w_{o,i}$ represents the weight of the i-th omics feature, $x_{N,i}$ is the i-th feature of the omics data for cell line N, $w_{p,k}$ is the weight of the k-th fingerprint feature, $f_{A,j}$ is the j-th feature of drug A's fingerprint, and c is a fitting constant.

The difference in sensitivity between drug A and drug B on the same cell line N is given by:

$$y_{A,N}-y_{B,N}=\sum_{k} w_{p,k}f_{A,j}-\sum_{k} w_{p,k}f_{B,j}=\sum_{k} w_{p,k}\left( f_{A,j}-f_{B,j} \right)$$

This difference is independent of the cell line N, indicating that the sensitivity difference between two drugs on any cell line is a constant. As such, the pairwise correlation between two drugs across cell lines will be 1, showing that a linear method is not suitable for predicting the sensitivity of novel drugs.
